# ProkEvo: an automated, reproducible, and scalable framework for high-throughput bacterial population genomics analyses

**DOI:** 10.1101/2020.10.13.336479

**Authors:** Natasha Pavlovikj, Joao Carlos Gomes-Neto, Jitender S. Deogun, Andrew K. Benson

## Abstract

Whole Genome Sequence (WGS) data from bacterial species is used for a variety of applications ranging from basic microbiological research, diagnostics, and epidemiological surveillance. The availability of WGS data from hundreds of thousands of individual isolates of individual microbial species poses a tremendous opportunity for discovery and hypothesis-generating research into ecology and evolution of these microorganisms. Scalability and user-friendliness of existing pipelines for population-scale inquiry, however, limit applications of systematic, population-scale approaches. Here, we present ProkEvo, an automated, scalable, and open-source framework for bacterial population genomics analyses using WGS data. ProkEvo was specifically developed to achieve the following goals: 1) Automation and scaling of complex combinations of computational analyses for many thousands of bacterial genomes from inputs of raw Illumina paired-end sequence reads; 2) Use of workflow management systems (WMS) such as Pegasus WMS to ensure reproducibility, scalability, modularity, fault-tolerance, and robust file management throughout the process; 3) Use of high-performance and high-throughput computational platforms; 4) Generation of hierarchical population-based genotypes at different scales of resolution based on combinations of multi-locus and Bayesian statistical approaches for classification; 5) Detection of antimicrobial resistance (AMR) genes, putative virulence factors, and plasmids from curated databases and association with genotypic classifications; and 6) Production of pan-genome annotations and data compilation that can be utilized for downstream analysis. The scalability of ProkEvo was measured with two datasets comprising significantly different numbers of input genomes (one with ~2,400 genomes, and the second with ~23,000 genomes). Depending on the dataset and the computational platform used, the running time of ProkEvo varied from ~3-26 days. ProkEvo can be used with virtually any bacterial species and the Pegasus WMS facilitates addition or removal of programs from the workflow or modification of options within them. All the dependencies of ProkEvo can be distributed via conda environment or Docker image. To demonstrate versatility of the ProkEvo platform, we performed population-based analyses from available genomes of three distinct pathogenic bacterial species as individual case studies (three serovars of *Salmonella enterica*, as well as *Campylobacter jejuni* and *Staphylococcus aureus*). The specific case studies used reproducible Python and R scripts documented in Jupyter Notebooks and collectively illustrate how hierarchical analyses of population structures, genotype frequencies, and distribution of specific gene functions can be used to generate novel hypotheses about the evolutionary history and ecological characteristics of specific populations of each pathogen. Collectively, our study shows that ProkEvo presents a viable option for scalable, automated analyses of bacterial populations with powerful applications for basic microbiology research, clinical microbiological diagnostics, and epidemiological surveillance.

## Introduction

Due to the advances in WGS technology, its decreasing costs, and the proliferation of publicly available tools and WGS-based datasets, the field of bacterial genomics is evolving rapidly from comparative analysis of a few representative strains of a given species, toward systematic, population-scale analyses of thousands genomes, which can provide new insights into their ecological characteristics and underlying evolutionary mechanisms [1,2,3,5]. The applications of WGS-based population genomics range from basic research, public health, pathogen surveillance, clinical diagnostics, and ecological and evolutionary studies of pathogenic and non-pathogenic species [3,4]. Indeed, use of WGS by public health agencies is providing unprecedented levels of resolution and accuracy and is becoming the standard for epidemiological surveillance, outbreak detection, and source-tracking [6,7,8].

Although the major applications of WGS-based genotyping in public health are focused on outbreak detection and source-tracking, the availability of large amounts of WGS data from populations of pathogenic bacteria from public health and regulatory agencies, and academic research creates tremendous opportunity for ecological and evolutionary inquiry at unprecedented scales of genomic resolution. For example, systematically monitoring the frequencies of specific genotypes of a pathogen, collected over time from the environment, food animals, and food production environments, can identify significant shifts in genotype frequencies that are driven by ecological events in the environment and/or within food production systems [10]. Powered statistically by the large number of genomes available from historical and ongoing surveillance, complex trait analyses can be used to identify causal variants and/or gene acquisition/loss associated with high-frequency vs. low-frequency genotypes, providing resourceful information that might explain the variation in ecological fitness across genotypes [9,10]. Genomic segments under different patterns of selection or associated with distinct populations based on serovars [13,14], or genotypes at different scales of resolution [15], can further be examined *in silico* to predict unique functional characteristics and phenotypes of populations (e.g. antimicrobial resistance (AMR) [11], virulence and metabolic attributes [10,12]), leading to empirically-determined hypotheses about the selective forces that are shaping these populations.

Currently, there are small number of automated pipelines available for analysis and genotypic classification of bacterial genomes, including EnteroBase [17], TORMES [18], Nullarbor [19], ASA3P [20]. These pipelines each have unique advantages, but differ in the programming language used, the size and type of supported input data, the supported bioinformatics tools, and the computational platform used. Consequently, these pipelines each support very different types of biological inquiry. Our work was motivated by the need for a scalable and portable, WGS-based population genomics platform that can accommodate scalable, hierarchical genotypic classifications and gene annotations for population-based inquiry (ecological, evolutionary, epidemiological). To accommodate the complex combinations of multiple, sequential data processing steps required for such a platform (where each step performs different task and requires different software), we used a powerful Workflow Management System (WMS) [21,22,23,24], that can efficiently manage massive numbers of computational operations in different types of high-performance computing environments, including University or publicly available clusters [25,26], clouds [27], or distributed grids [28,29].

In this paper, we describe ProkEvo – an automated and user-friendly platform for population-based inquiry of bacterial species that is managed through the Pegasus WMS and is portable to computing clusters, clouds, and distributed grids. ProkEvo works with raw paired-ended Illumina reads as input, and is composed of multiple sequential steps for processing and analysis of data from hundreds to tens of thousands of genomes. For each input genome, these steps include trimming and quality control, genome assembly, serovar prediction (in the case of *Salmonella enterica*), genotypic classification based on multilocus-sequence typing (MLST) using seven or approximately 300 loci, genotypic classification based on heuristic Bayesian genomic admixture analysis at different scales of resolution, screening for AMR and putative virulence genes, plasmid identification, and pan-genome analysis.

Here, we show the utility and adaptability of ProkEvo for basic metrics of population genetic analysis of three serovars of the enteric pathogen *S. enterica* (serovars Typhimurium, Newport and Infantis), as well as the pathogens *C. jejuni* and *S. aureus*. We also demonstrate the scalability and modularity of ProkEvo with datasets having ~2,400 and ~23,000 genomes and further illustrate the portability and performance of ProkEvo on two different computational platforms, the University of Nebraska high-performance computing cluster (Crane) and the Open Science Grid (OSG), a distributed, high-throughput cluster. Because of the multi-disciplinary environments required for implementation and applications of ProkEvo, we also provide initial guidance for researchers on utilization of some of the output files generated by ProkEvo to perform meaningful population-based analyses in a reproducible fashion using a combination of R and Python scripts.

## Materials & Methods

### Overview of ProkEvo

The ProkEvo pipeline is capable of processing raw, paired-end Illumina reads obtained from tens of thousands of genomes present in the NCBI database utilizing high-performance and high-throughput computational resources. The pipeline is composed of two sub-pipelines: 1) The first sub-pipeline performs the standard data processing steps of sequence trimming, *de novo* assembly, and quality control; 2) The second sub-pipeline uses the assemblies that have passed the quality control and performs specific population-based classifications (serotype prediction specifically for Salmonella, genotype classification at different scales of resolution, analysis of core- and pan-genomic content). Pegasus WMS manages and splits each sub-workflow into as many independent tasks as possible to take advantage of many computational resources.

A text file of SRA identifications corresponding to raw Illumina reads available from the Sequence Read Archive (SRA) database in NCBI (NCBI SRA) is used as an input to the pipeline. The first step of the pipeline and the first sub-workflow is automated download of genome data from NCBI SRA [30]. This is done using the package parallel-fastq-dump [31]. The SRA files are downloaded using the prefetch utility, and the downloaded files are converted into paired-end fastq reads using the program parallel-fastq-dump. While the SRA Toolkit [30] provides the same functionality, this toolkit can be slow sometimes and show intermittent timeout errors, especially when downloading many files. parallel-fastq-dump is a wrapper for SRA Toolkit that speeds the process by dividing the conversion to fastq files into multiple threads. After the raw paired-end fastq files are generated, quality trimming and adapter clipping is performed using Trimmomatic [32]. FastQC is used to check and verify the quality of the trimmed reads [33] and it is run independently for each paired-end dataset with concatenation of all output files at the end for a summary. The paired-end reads are assembled *de novo* into contigs using SPAdes [34]. These assemblies are generated using the default parameters. The quality of the assemblies is evaluated using QUAST [35]. The information obtained from QUAST is used to discard assemblies with 0 or more than 300 contigs, or assemblies with N50 value of less than 25,000. QUAST-based filtering of the assemblies concludes the first part or first sub-pipeline of the workflow. Each of these steps is independent of the input data and each task is performed on one set of paired-end reads using one computing core. This makes the analyses modular and suitable for high-throughput resources with many available cores. Moreover, having many independent tasks significantly reduces the memory and time requirements while generating the same results as when the analyses are done sequentially. Thus, if a dataset has paired-end reads from 2,000 different genomes and a computational platform has 2,000 available cores, ProkEvo will scale and utilize all these resources at the same time.

The second sub-pipeline uses the assemblies which passed quality control to perform specific population-based characterizations, including genotypic classifications, serovar prediction (exclusively for Salmonella), gene-based annotations, and pan-genome outputs. PlasmidFinder is used to identify plasmids in the assemblies [36]. PlasmidFinder comes with curated database of plasmid replicons to identify plasmids in the WGS data (currently enriched for plasmids from the Enterobacteriaceae). SISTR is used for Salmonella and produces serovar prediction and *in silico* molecular typing by determination of antigen gene and core-genome multilocus-sequence typing (cgMLST) gene alleles [14]. SISTR generates multiple output files. Of primary interest for downstream analyses is the main SISTR output file named sistr_output.csv. The filtered assemblies are annotated using Prokka [37], which is based on a curated set of core and HMM databases for the most common bacterial species. If needed, one can customize and create their own annotation database. In addition to the other files, Prokka produces annotation files in GFF3 format that are used with Roary [38] to identify the pan-genome and to generate core-genome alignments. The core-genome alignment file produced is then used with fastbaps, an improved version of the BAPS clustering method [39], to hierarchically cluster the genomic sequences from the multiple sequence alignment in varying numbers of stratum (i.e. levels of resolution). Multilocus-sequence typing is also performed on the assemblies using MLST [40]. Here, the filtered genome assemblies from individual bacterial isolates are categorized into specific genotypes based on allele combinations from seven ubiquitous, house-keeping genes [41]. In addition to these analyses, the filtered assemblies are screened for AMR and virulence associated genes using ABRicate [42]. ABRicate comes with multiple comprehensive gene-based mapping databases, and the ones used in ProkEvo are NCBI [43], CARD [44], ARG_ANNOT [45], Resfinder [46], and VFDB [47]. Prokka, SISTR, PlasmidFinder, MLST, and ABRicate are independent of each other, and they are all run simultaneously in parallel. Moreover, Prokka, SISTR and PlasmidFinder perform their computations per filtered assembly, while MLST and ABRicate require all filtered assemblies to be used together. Running multiple independent jobs simultaneously is one of the key factors to maximize computational efficiency. With respect to Salmonella genomes, once the SISTR analyses finish for all assemblies, the generated independent sistr_output.csv files are concatenated. This aggregation of files can be done because the genome categorization to serovars and cgMLST lineages done by SISTR occurs completely independent for each genome. Each tool executed in ProkEvo is run with specific options set as defaults. While the options used in this paper fit the presented case studies, these options are easily adjustable and configurable in the pipeline. Because we developed ProkEvo for studying a diverse array of bacterial species, the pipeline was specifically designed to incorporate programs such as SISTR for *Salmonella enterica*, were serovar classifications can be made accurately based on the Kauffman-White scheme [56]. However, other serotype prediction modules can be substituted for SISTR to accommodate user-specific needs. Additionally, MLST program can be directed to species-specific sets of genetic loci used for classification, as shown with the *Campylobacter jejuni* and *Staphylococcus aureus* datasets.

The modularity of ProkEvo allows us to decompose the analyses into multiple tasks, some of which can be run in parallel, and utilize a WMS. ProkEvo is dependent on many well-developed bioinformatics tools and databases, the setup and installation of which are not trivial. To make this process easier, reduce the technical complexity, and allow reproducibility, we provide two software distributions for ProkEvo. The first distribution is a conda environment that contains all software dependencies [48], and the second is a Docker image that can be used with Singularity [49]. Both distributions are supported by the majority of computational platforms and integrate well with ProkEvo, and can be easily modified to include other tools and steps. The code for ProkEvo, and both the conda environment and the Docker image, are publicly available at our GitHub repository (https://github.com/npavlovikj/ProkEvo).

### Pegasus Workflow Management System

ProkEvo uses the Pegasus WMS, which is a framework that automatically translates abstract, high-level workflow descriptions into concrete efficient scientific workflows that can be executed on different computational platforms such as clusters, grids, and clouds. The abstract workflow of Pegasus WMS contains information and description of all executable files (transformations) and logical names of the input files used by the workflow. Complementing the abstract component is a concrete workflow, which specifies the location of the data and the execution platform [24]. The workflow is organized as a directed acyclic graph (DAG), where the nodes are the tasks and the edges are the dependencies. Next, the workflow is submitted using HTCondor [50]. Pegasus WMS uses DAX (directed acyclic graph in XML) files to describe an abstract workflow. These files can be generated using programming languages such as Java, Perl, or Python. The high-level of abstraction of Pegasus allows users to ignore low-level configurations required by the underlying execution platforms. Pegasus WMS is an advanced system that supports data management and task execution in automated, reliable, efficient, and scalable manner. This whole process is monitored, and the workflow data is tracked and staged. The requested output results are presented to the users, while all intermediate data can be removed or re-used. In case of errors, jobs are automatically re-initiated. If the errors persist, a checkpoint file is produced so the job can be resubmitted and resumed. Pegasus WMS supports sub-workflows, task clustering and defining memory and time resources per task.

Pegasus WMS also generates web dashboard for each workflow for better workflow monitoring, debugging, and analyzing, which helps users to analyze workflows based on useful statistics and metrics of the workflow performance, running time, and machines used. ProkEvo uses Python to create the workflow description. Each step of the pipeline is a computational job represented as a node in the DAG. Two nodes are connected with an edge if the two jobs need to be run one after another. The input and output files are defined in the DAG as well. All jobs that are not dependent on each other can be run concurrently. Each job uses its own predefined script that executes the program required by the job with the specified options. This script can be written in any programming language. The bioinformatics tools and programs required by ProkEvo can be distributed through conda environment [48] or Docker image [49]. The predefined scripts within this release of ProkEvo enable running without further change or modification. With the modularity of Pegasus, each job requests its own run time and memory resources. Exceeding the memory resources is a common occurrence in any bioinformatics analysis and based on this assumption, when exceeding the memory is a reason for a job failure, Pegasus retries the job with increased requirements. Higher memory requirements may mean longer waiting times for resources, and the Pegasus WMS uses high memory resources only when needed. ProkEvo is developed in a way that supports execution on various high-performance and high-throughput computational platforms. In the analyses for this paper, we use both the University cluster and OSG, and working versions for both platforms are available in our GitHub repository (https://github.com/npavlovikj/ProkEvo).

### Computational execution platforms

Traditionally, data-intensive scientific workflows have been executed on high-performance and high-throughput computational platforms. While high-performance platforms provide resources for analyses that require significant numbers of cores, time, and memory, high-throughput platforms are suitable for many small and short independent tasks. The design of ProkEvo is suitable for different computational environments like University and other publicly or privately available clusters and grids, and thus provides flexibility in the computational platform. We have evaluated ProkEvo on two different computational platforms—a University cluster and the distributed Open Science Grid.

### University cluster (Crane), a high-performance computational platform

University and other public clusters are shared by diverse communities of users and enforce fair-share scheduling and file and disk spaces quotas. These clusters are suitable for various types of jobs, such as serial, parallel, GPU, and high memory specific jobs, thus the high-performance.

Crane [25] is one of the high-performance computing clusters at the University of Nebraska Holland Computing Center (HCC). Crane is Linux cluster, having 548 Intel Xeon nodes with RAM ranging from 64GB to 1.5TB, and it supports Slurm and HTCondor as job schedulers. In order to use Crane, users obtain an HCC account associated with a University of Nebraska faculty or research group. Importantly, most University and publicly available high-performance clusters are administered in a manner similar to Crane and would be suitable for running ProkEvo.

Crane has support for Pegasus and HTCondor, and no further installation is needed in order to run ProkEvo. Due to the limited resources and fair-share policy on Crane, tens to hundreds of independent jobs can be run concurrently. We provide a version of ProkEvo suitable for Crane with conda environment, which contains all required software. Crane has a shared file system where the data is accessible across all computing nodes. Depending on the supported file system, Pegasus is configured separately and handles the data staging and transfer accordingly. However, users do not need advanced experience in high-performance computing to run ProkEvo on Crane, or most other University or publicly available clusters. Users only need to provide list of SRA identifications and run the submit script that distributes the jobs automatically as given in our GitHub repository (https://github.com/npavlovikj/ProkEvo).

### Open Science Grid (OSG), a distributed, high-throughput computational platform

The Open Science Grid (OSG) is a distributed, high-throughput distributed computational platform for large-scale scientific research [28,29]. OSG is a national consortium of more than 100 academic institutions and laboratories that provide storage and tens of thousands of resources to OSG users. These sites share their idle resources via OSG for opportunistic usage. Because of its opportunistic approach, OSG as a platform is ideal for running massive numbers of independent jobs that require less than 10GB of RAM, less than 10GB of storage, and less than 24 hours running time. If these conditions are fulfilled, in general, OSG can provide unlimited resources with the possibility of having hundreds or even tens of thousands of jobs running at the same time. The OSG resources are Linux-based, and due to the different sites involved, the hardware specifications of the resources are different and vary. Access and use of OSG is free for academic purposes and the user’s institution does not need to be part of OSG to use this platform.

All steps from the population genomics analyses of ProkEvo fulfill the conditions for OSG-friendly jobs and ProkEvo can efficiently utilize these distributed high-throughput resources to run thousands of analyses concurrently when the resources are available. OSG supports Pegasus and HTCondor, so no installation steps are required. We provide version of ProkEvo suitable for OSG (https://github.com/npavlovikj/ProkEvo). This version uses the Docker image with all software requirements via Singularity and supports non-shared file system. In non-shared systems, the resources do not share the data. The data are read and written from a staging location, all of which is managed by the Pegasus WMS. In order to run ProkEvo on OSG, users only need to provide list of SRA identifications and run the submit script without any advanced experience in high-throughput computing.

### Population genomics analyses

The population-based analyses performed in this paper provide an initial guidance on how to comprehensively utilize the following output files produced by ProkEvo: 1) MLST output (.csv) [40]; 2) SISTR output (.csv) [14]; 3) BAPS output (.csv) [39]; 4) Core-genome alignment file (core_gene_alignment.aln) for phylogenetic analysis [38]; and 5) Resfinder output (.csv) containing AMR genes [46]. We use both R and Python 3 Jupyter Notebooks for all our data analyses (https://github.com/npavlovikj/ProkEvo). The input data used for these analyses is available on Figshare (https://figshare.com/projects/ProkEvo/78612).

A first general step in this type of analysis is opening all files in the preferred environment (i.e. RStudio or JupyterHub), and merging them into a single data frame based on the SRA (genome) identification. Next, we perform quality control (QC) of the data, focusing on identifying and dealing with missing values, or cells of the data frame containing erroneous characters such as hyphens (-) and interrogation marks (?). For that, we demonstrate our approach for cleaning up the data prior to conducting exploratory statistical analysis and generating all visualizations.

In the case of Salmonella datasets, an additional “checking/filtering” step was used after the QC is complete. Since the program SISTR provides a serovar call based on genotypic information, one can opt for keeping or excluding those genomes that do not match the original serovar identification in the analysis. Both approaches are justifiable with the latter one being more conservative, and it specifically assumes that the discordance between data entered in NCBI and genotypic prediction done by SISTR is accurate. However, it is important to remember that we initially expect that the dataset belongs to a particular serovar because of the keywords we used to search the NCBI SRA database, such as: “*Salmonella* Newport”, “*Salmonella* Typhimurium”, or “*Salmonella* Infantis”. Typically, the proportion of genomes that are classified differently by SISTR than the designation associated with the file in SRA is ~<3% for any given Salmonella dataset tested here. In our application, we chose a conservative approach and either filtered the “miscalls” out of the data, or kept it as a separate group called “other serovars”. The latter approach was done for specific analyses, such as phylogenetics, where the program required the use of all data points in place (e.g. ggtree in R). This situation arises because the core-genome alignment used for the phylogeny is generated by Roary without considering the SISTR prediction for serovar calls. If such consideration is relevant, the user can add a condition to the pipeline to run Roary after considering SISTR results, but this situation only applies to Salmonella genomes. However, we do note that stringent requirements for serotype classification (i.e. filtering out “miscalls” based on SISTR predictions) could eliminate important variants that may genotypically match known populations of the serovar, but which have acquired mutations or recombination events at serotype-determining loci. Our suggestion is that for any predictive analysis, one should either filter out, or at least, classify the potential miscalls as other serovars after running SISTR.

To define hierarchical relationships of genotypic classifications at varying levels of resolution, the ProkEvo pipeline combines multi-locus MLST-based genotypes at different scales of resolution with Bayesian-based admixture analysis (BAPS), which classifies genomes based on core-genomic structure (i.e. only shared content). The BAPS-based approach to genomic classifications is callable, and allows the user to circumvent computationally-intensive use of phylogeny, which is not scalable to thousands of core genomes. Thus, evolutionary “familial” relationships across STs or thousands of cgMLST genotypes can be inferred by their hierarchical relationships to BAPS-based classifications. In this version of ProkEvo, we have implemented legacy MLST for ST calls using seven loci, core-genome MLST (cgMLST) that uses approximately 330 loci for MLST analysis, and a Bayesian-based BAPS haplotype classification using six layers of BAPS (BAPS1 being the lowest level of resolution and BAPS6 being the highest). To explore the hierarchical relationships of genotypes, one can simply examine the distribution of legacy STs among genomes having similar admixtures based on classification at the lowest level of BAPS resolution (BAPS1). Likewise, the genetic relationships of thousands of cgMLST genotypes can also be assessed with respect to the BAPS-based and ST-linked genomic architecture at different levels of BAPS resolution to infer evolutionary relationships.

The hierarchical approach was possible for the *S.* Newport dataset of ~2,400 genomes (USA data), but the core-genome alignment step was not scalable to the 10-fold larger dataset of *S.* Typhimurium (~23,000 genomes – worldwide data). This larger dataset was split into twenty smaller datasets during the core-genome alignment step. Although random partitioning of the subsets should yield the same classifications of dominant genomic groups, the Bayesian classification algorithm (BAPS) will not necessarily assign the same genomic types in different datasets to the same group numbers. Thus, aggregation of the BAPS data from multiple, independently analyzed subsets requires user-based input. On the other hand, sub-setting larger datasets is advantageous for downstream data science and machine learning analyses since they require a nested cross-validation approach for feature selection and predictive analytics. Herein, we used a random sampling approach to create subsets of the genomic data for the large number of *S.* Typhimurium genomes that were input into Roary. Based on the number of genomes, we created 20 subsets, each having 1,076-1,077 genomes. Next, from the GFF files produced by Prokka, we randomly selected and assigned genomes to each group using custom Bash scripts. Both Roary and fastbaps were run per group, resulting in 20 independent runs with the corresponding output files. To evaluate randomness of subset assignments, haplotype frequencies of the major cgMLST genotypes are assessed for each subset at each of the six levels of BAPS-based resolution (BAPS1-6). A highly clonal population of multiple cgMLST genotypes would be expected to group into a single admixture-based BAPS genomic type at the lowest level of BAPS1 resolution and remain confined to one or a small number of BAPS genomic types at increasing levels of BAPS-based resolution. In contrast, a diverse population of cgMLSTs that are more distantly related (e.g. not highly clonally related) will partition between multiple BAPS-based genomic groups. In practice, this analysis is important for examining the degrees of population heterogeneity and diversification, which has implications for ecological and epidemiological inference.

Complementary to this population structure analysis, we also measured distributions of AMR genes within and between Salmonella serovars, including *S*. Infantis (~1,700 genomes – USA data). Within serovar, the frequencies of AMR genes were estimated between major ST genotypes using the Resfinder outputs for identification of putative AMR genes. We arbitrarily selected genes with proportion higher than or equal to 25% for *S*. Newport, *S*. Infantis, and *S*. Typhimurium, for visualizations, which were produced with ggplot2 in R [52]. The respective scripts are provided in our repository (see GitHub link for code).

To demonstrate the versatility of ProkEvo across multiple species, we also conducted a population-based analysis of *C. jejuni* and *S. aureus* datasets comprising isolates from the USA, containing 21,919 and 11,990 genomes, respectively. For both datasets, we analyzed the population structure using BAPS1 and STs. The same hierarchical population basis described for Salmonella applies here, with BAPS1 coming first and STs next in terms of population ranking. We used a random sample of ~1,000 genomes of each species to demonstrate the distribution of BAPS1 and STs onto the phylogenetic structure. Phylogenies were constructed using the core-genome alignment produced by Roary, and by applying the FastTree program [53] using the generalized time-reversible (GTR) model of nucleotide evolution without removing genomic regions putatively affected by recombination (see GitHub link for code). Additionally, we showed the distribution of STs within each bacterial species (only showed STs with proportion higher than 1%), and the relationship between the relative frequencies of dominant STs and AMR genes. Genes with relative frequency below 25% were filtered out of the data. All visualizations were generated with ggplot2 in R, and the scripts are also provided in our repository.

## Results

### Overview of ProkEvo

The overall flow of tasks performed in ProkEvo is illustrated in *Fig. 1*, including all specific bioinformatics tools used for each task. The DAG shown in *Fig. 2* represents the Pegasus WMS design of ProkEvo and it shows all independent input and output files, tasks, and the dependencies among them. The modularity of ProkEvo allows every single task to be executed independently on a single core. As seen on *Fig. 2*, there are approximately 10 tasks executed per genome. When ProkEvo is used with whole bacterial populations of thousands of genomes, the number of total tasks is immense.

**Figure 1:**
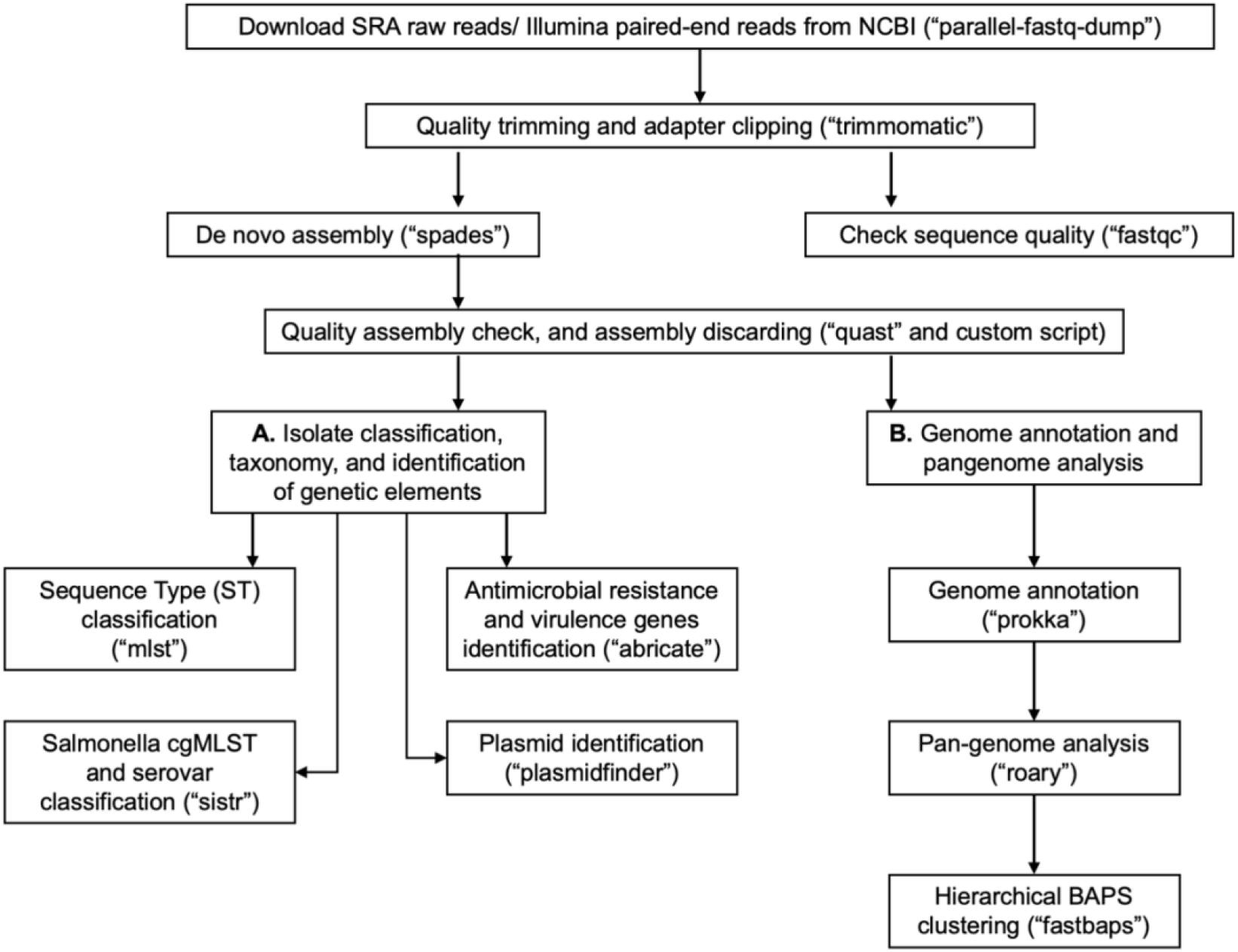
Overall ProkEvo’s computational workflow. Top-down flow of tasks for the ProkEvo pipeline. The squares represent the steps, where the bioinformatics tool used for each step is shown in brackets. The pipeline starts with downloading raw Illumina sequences from NCBI, after providing a list of SRA identifications, and subsequently performing quality control. Next, *de novo* assembly is performed on each genome using SPAdes and the low-quality contigs are removed. This concludes the first part of the pipeline, the first sub-workflow. The second sub-workflow is composed of more specific population-genomics analyses, such as genome annotation and pan-genome analyses (with Prokka and Roary), isolate cgMLST classification and serotype predictions from genotypes in the case of Salmonella (SISTR), ST classification using the MLST scheme, non-supervised heuristic Bayesian genotyping approach using core-genome alignment (fastbaps), and identifications of genetic elements with ABRicate and PlasmidFinder.

**Figure 2:**
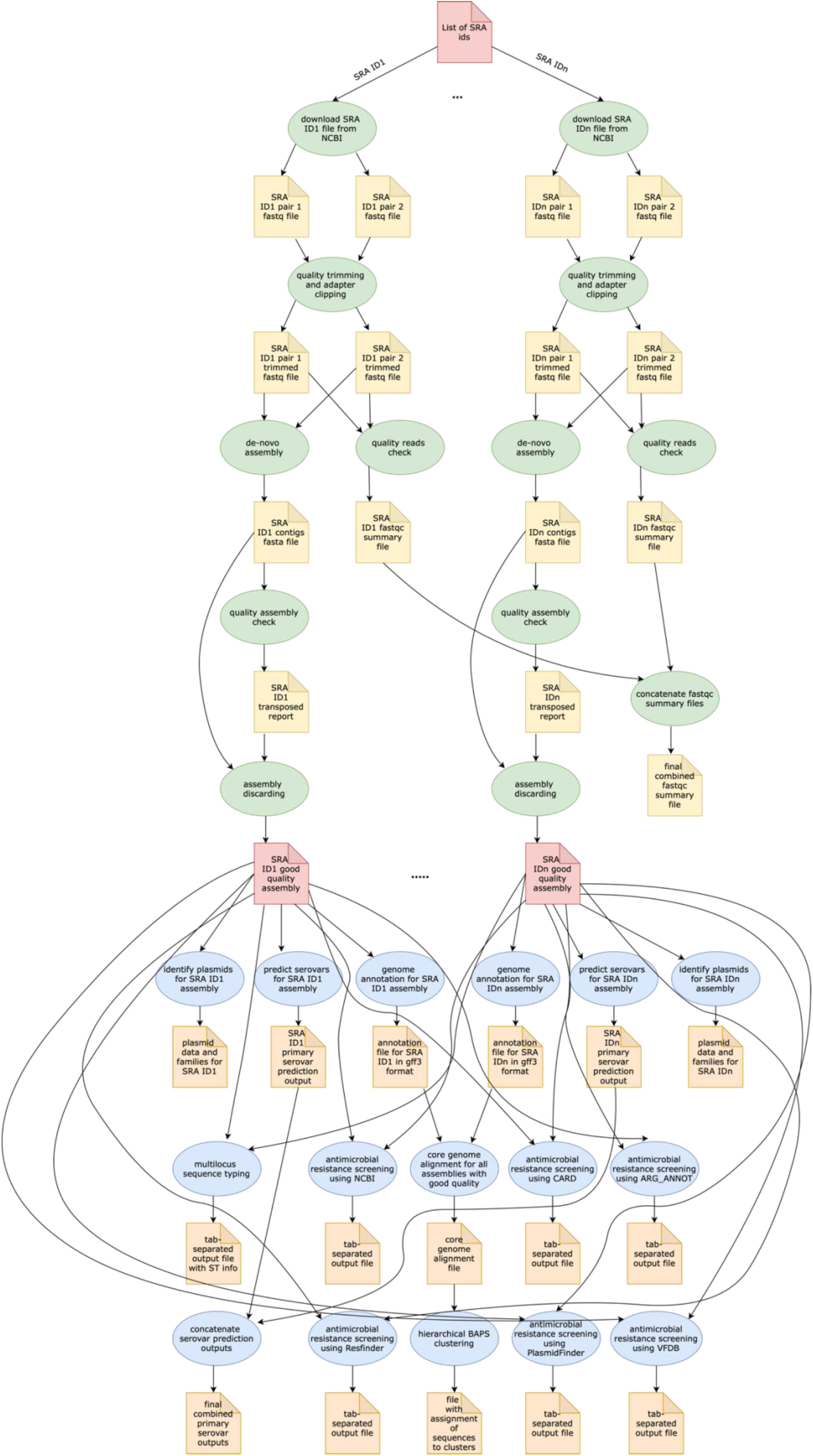
Pegasus workflow of ProkEvo. Pentagons represent the input and output files, the ovals represent the tasks (jobs), and the arrows represent the dependency order among the tasks. Pentagons are colored in red for the input files used for the first and second sub-workflow, respectively. The yellow pentagons and the green ovals represent the input and output files, and tasks (jobs) that are part of the first sub-workflow. The pentagons colored in orange and the ovals colored in blue are the input and output files, and tasks used in the second sub-workflow. While the first sub-workflow is more modular, most of the tasks from the second sub-workflow are performed on all processed genomes together. Here, the steps of the analyses for two genomes are shown, and those steps and tasks remain the same regardless of the number of genomes. The number of tasks significantly increases with the number of genomes used, and because of the modularity of ProkEvo, each task is run on a single core which facilitates parallelization at large scale. Theoretically, if there are *n* cores available on the computational platform, ProkEvo can utilize all of them and run the corresponding *n* independent tasks, simultaneously (1:1 correspondence).

To evaluate capability of the Pegasus WMS to scale tasks independently on diverse computational platforms, ProkEvo was run with two datasets of significantly different size (~2,400 genomes [1X] vs. ~23,000 genomes [10X]) on two different computational platforms— the University of Nebraska high-performance computing cluster (Crane) and the Open Science Grid (OSG), a distributed, high-throughput cluster (*Fig. 3*). The ProkEvo code available on our GitHub page supports both platforms and each platform has unique structure and idiosyncratic advantages and disadvantages (*Fig. 3*). Each dataset was run once on the two platforms and performance metrics were collected for the Pegasus WMS workflow. Of note, there may be variation in the ProkEvo runtime from project to project based on the availability of resources on each platform. As an HPC resource of the Holland Computing Center, the Crane cluster is managed by fair-share scheduling, while as an opportunistic HTC resource, the OSG resources may be dynamically de-provisioned or having intermittent issues. These factors may impact the future predictability of running time and performance of ProkEvo on both platforms. On average, we had hundreds of jobs running at a time on Crane, and because of the similar type of nodes available, the runtime should be similar for multiple runs of the same workflow. On the other hand, the nodes on OSG are more diverse and the runtime and the number of jobs for multiple runs can be significantly different (from few jobs running at the same time to few tens of thousand).

**Figure 3:**
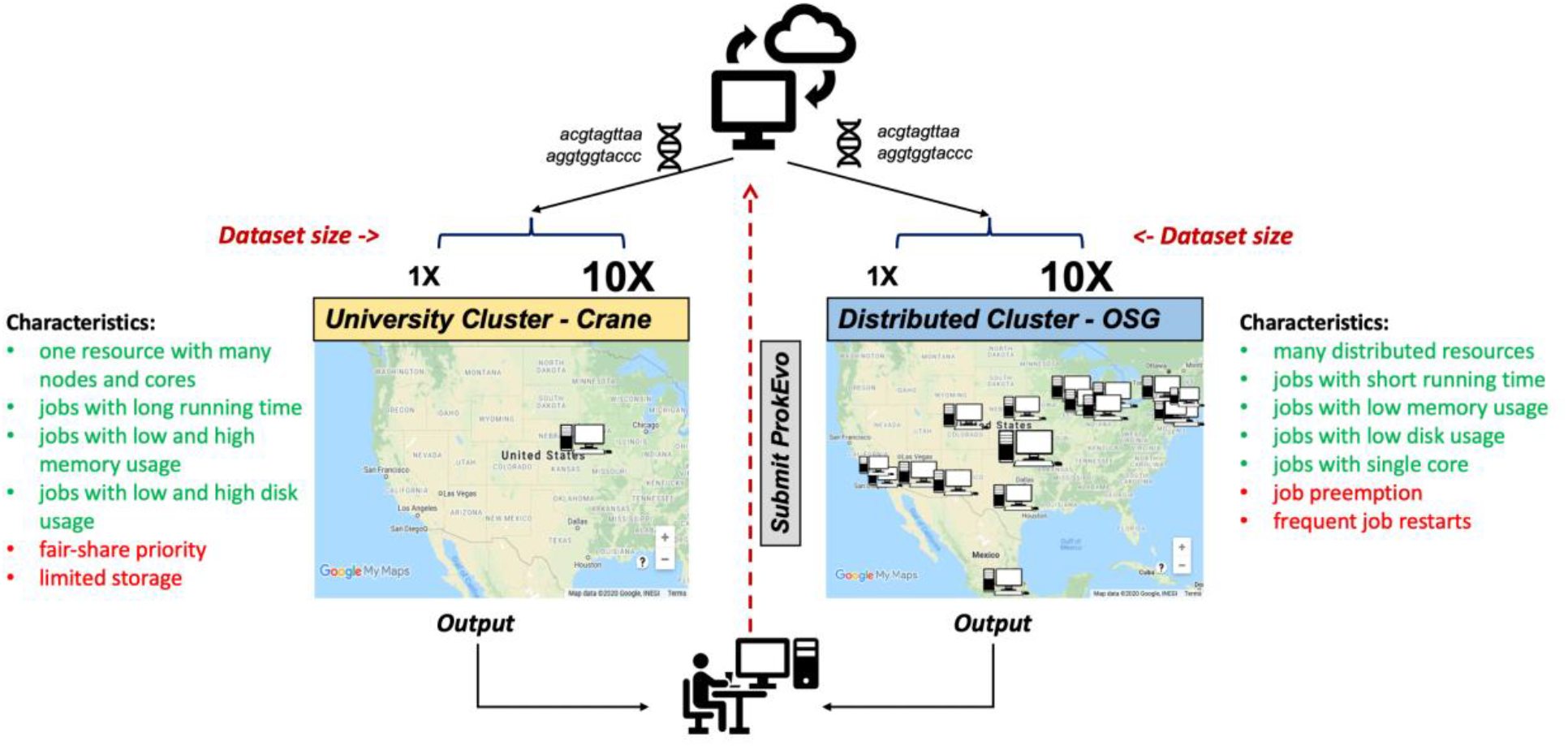
Computational experimental approach to test the performance of ProkEvo using two different computational platforms with datasets of different size. To test how ProkEvo would perform with a small (1X) vs. moderately large (10X) datasets, in addition to using different computational resources, we have designed the following experiment: 1) Selected two adequately sized datasets including genomes from *S.* Newport (1X – from USA) and *S.* Typhimurium (10X – worldwide); 2) Used two different types of computational platforms: Crane, the University of Nebraska high-performance computing cluster, and the Open Science Grid, as a distributed high-throughput computing cluster; 3) We then ran both datasets on the two platforms with ProkEvo, and collected the statistics for the performance in order to provide a comparison between the two different computational platforms, as well as possible guidance for future runs. Of note, the text in green and red correspond to advantages and disadvantages of using each computational platform, respectively. Map data ©2020 Google, INEGI.

ProkEvo consists of two sub-workflows, with number of jobs varying from a few thousands to a few hundreds of thousands, depending on the dataset. “pegasus-statistics” generates summary metrics/statistics regarding the workflow performance, such as the total number of jobs, total run time, number of jobs that failed and succeeded, task and facility information, etc. The total distributed running time is the total running time of ProkEvo from the start of the workflow to its completion. The total sequential running time is the total running time if all steps in ProkEvo are executed one after another. In case of retries, the running times of all re-attempted jobs are included in these statistics as well. Beside the workflow runtime information, *Table 1* also shows the maximum total number of independent jobs ran on Crane and OSG within one day. Moreover, the total count of succeeded jobs is shown for both computational platforms and datasets.

**Table 1:**
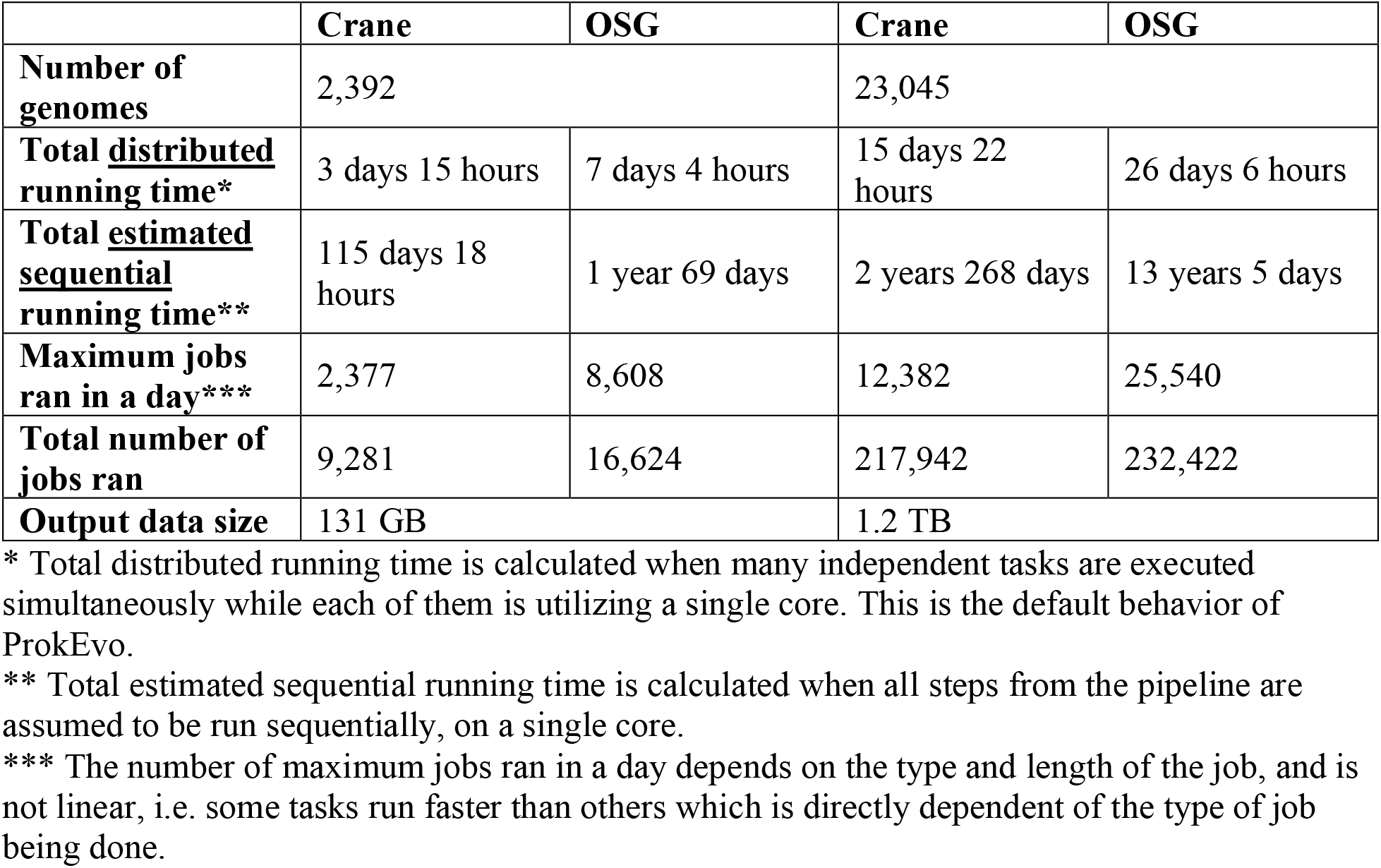
Comparison of ProkEvo’s performance on Crane and OSG with two datasets with significant difference in size and number of genomes.

When ran on Crane, ProkEvo with *S.* Newport completely finished in 3 days and 15 hours. If this workflow were run sequentially on Crane, its cumulative running time would be 115 days and 18 hours. On the other hand, ProkEvo with *S.* Newport finished in 7 days and 4 hours when OSG was used as a computational platform. Similarly, if this workflow were run sequentially on OSG, its cumulative running time would be 1 year and 69 days. As it can be observed, the workflow running on OSG took longer than the workflow running on Crane. OSG provides variable resources with different configuration and hardware, and depending on that, the performance may vary significantly. Also, the OSG jobs may be preempted if the resource owner submits more jobs. In this case, the preempted job is retried, but that additional time is added to the workflow wall time. While the maximum number of independent jobs ran on Crane in one day is 2,377, this number is 8,606 when OSG was used. This is where the importance of using HTC resources such as OSG can be observed - the high number of jobs and nodes that can be run and used simultaneously, which is often a limit for University clusters. The total number of successful jobs ran with ProkEvo for the *S.* Newport dataset is 9,281 on Crane and 16,624 on OSG. Due to the opportunistic nature of the OSG resources, a running job can be cancelled and retried again, thus the higher number of jobs reported by OSG. Similar pattern can be observed when ProkEvo was run with the *S.* Typhimurium dataset. When ran on Crane, ProkEvo with *S.* Typhimurium completely finished in 15 days and 22 hours. If this workflow was run sequentially on Crane, its cumulative running time would be 2 years and 268 hours. On the other hand, the ProkEvo run for *S.* Typhimurium finished in 26 days and 6 hours, when using OSG as a computational platform. Similarly, if this workflow were run sequentially on OSG, its cumulative running time would be 13 years and 50 days. The maximum number of independent jobs ran on Crane and OSG is 12,382 and 25,540 respectively. The total number of successful jobs ran with the *S.* Typhimurium dataset is 217,942 on Crane and 232,422 on OSG.

Although the workflow run time was better when Crane was used as a computational platform, it should be pointed out that OSG is more efficient for datasets where there are more jobs running and, in our case, more genomes analyzed. As long as resources are available and no preemption occurs, workflows running on OSG can have a great performance. On OSG, ProkEvo ran on resources shared by thirty-four different facilities. Failures and retries are expected to occur on OSG, and their proportion may vary. From our experience, the number of failures and retries took up ~0.3%-30% of the total number of jobs. However, the OSG support staff acts promptly on isolating these issues, which can also be masked by a resilient and fault-tolerant workflow management systems like Pegasus WMS. All the data, intermediate and final files generated by ProkEvo are stored under the researcher’s allocated space on the file system on Crane. Depending on the file system, it is possible that there are file count and disk space quotas. When large ProkEvo workflows are run, these quotas may be exceeded. On the other hand, due to the non-shared nature of the file system of OSG, intermediate files are stored on different sites, and exceeding the quotas is usually not an issue.

Both Crane and OSG are computational platforms that have different structure and target different type of scientific computation. All analyses performed with ProkEvo fit both platforms well. Thus, we provide an unambiguous comparison of both platforms and show their advantages and drawbacks when large-scale workflows such as ProkEvo are run.

### Applications

To demonstrate various applications of ProkEvo to population genomic analysis of different bacterial pathogens, we used publicly available datasets from three phylogenetically distinct species of pathogens, including zoonotic serovars of *Salmonella*, as well as *C. jejuni* and *S. aureus*. While these datasets are clearly biased by inflation with clinical isolates and under-sampling from other environments (e.g. animal or environmental), our analysis emphasizes the utilities, approaches, and applications that can result from formal population-based analysis with ProkEvo as opposed to formal hypothesis-testing on the biased datasets. To achieve our objective, we present a series of independent case studies that encapsulate some of the most generally useful approaches for studying bacterial populations. Some important background concepts regarding bacterial population genetics and biology of each of these pathogens are described below. Notably, ProkEvo can be used with essentially any bacterial species with a few limitations: 1) The MLST program only works if the target bacterial species has an allelic profile present in the database, or is incorporated by the user; and 2) SISTR is designed specifically for analysis of Salmonella, but the program can be easily blocked out from the pipeline by the user.

### Overview of the population structure and ecology for *Salmonella enterica*, *Campylobacter jejuni*and *Staphylococcus aureus*

To understand utilities of ProkEvo from the case studies presented below, it is important that users are familiar with relevant aspects of the biology of the target organisms. In this report, we focus on three different species of foodborne pathogens, *Salmonella enterica*, *Campylobacter jejuni*, and *Staphylococcus aureus*, that are common worldwide [54] but are evolutionarily quite distinct from one another and have unique aspects of their biology. Foodborne gastroenteritis is among the most prevalent zoonotic infectious illnesses of humans, with *S. enterica* and *C. jejuni* being two of the most common etiologic bacterial agents. Foodborne illnesses can also be caused by production of toxins as organisms grow in the food matrix itself (intoxication) and *S. aureus* is among the most prevalent causative agents of food intoxication. However, due to the lack of reliable metadata we were not able to differentiate which *S. aureus* genomes are associated with foodborne illnesses, skin infections, or any other type of disease. Nonetheless, our goal here is to show the plasticity of ProkEvo in terms of being readily usable to different bacterial species.

The genus *Salmonella* is a member of the Phylum Proteobacteria and populations of these organisms can be found as common inhabitants of the gastrointestinal tract in a wide range of mammals, birds, reptiles, and insects and these organisms are often transmitted to humans through contaminated animal products, vegetables, fruits, and processed foods [55]. The genus *Salmonella* comprises two primary species (*S. enterica* and *S. bongori*), which are believed to have diverged from their last common ancestor approximately 40 million years ago [93]. Worldwide, *S. enterica* is the most frequently isolated species from human clinical cases and from most environments. The extant populations evolving through the *S. enterica* lineage have further diversified into six different sub-species. The vast majority (>90%) of known human cases are caused by populations descending from a single sub-species, namely *S. enterica subsp.* enterica (lineage I). Within lineage I *S.* enterica, there is still tremendous genetic and phenotypic diversity, which has historically been recognized by the diverse array of distinct sub-populations differentiated serologically as “serovars” by the combinations of lipopolysaccharide molecules and major protein components of their flagella on their cell surfaces (the Kauffman-White scheme) [56,94]. More than 2,500 serovars have been defined in *Salmonella enterica*. Serovars represent relevant biological units for epidemiological surveillance and tracking, because isolates belonging to the same serovar show much less variation with respect to important traits such as range of host species, survival in the environment, efficiency of transmission to humans, and virulence characteristics, than isolates from different serovars [7,94]. Moreover, Kaufmann-White-based serotypes are covariates with the population structure of *S. enterica* lineage I, with most serovars being found exclusively within a unique STs or ST clonal complexes, and consequently, the serotype of most isolates can be predicted accurately from MLST-based STs [94]. Here, we used genomes representing three serovars of *S. enterica* lineage I: *S*. Infantis, *S*. Newport, and *S*. Typhimurium. These are among the top twenty-five most prevalent and important zoonotic serovars of *Salmonella* according to the Center for Disease Control and Prevention [57], but have distinct ecologies. All three serovars are capable of gastroenteritis in humans and have reservoirs in livestock. Bovine appears to be the most common source for *S*. Infantis and *S*. Newport, while *S.* Typhimurium has a generalist life-style and can be found in swine, poultry, bovine, etc. [55].

The population structure of *Salmonella* is largely clonal and hierarchical genotyping schemes such as MLST show that isolates having genetically-related core-genome MLST (cgMLST) genotypes (high-resolution based on ~330 highly conserved genes) are mostly embedded within clonally-related STs defined at lower resolution by seven-gene MLST [7]. Thus, the *S. enterica* lineage I population structure can be hierarchically analyzed by first classifying based on the serovar, and then increasing levels of resolution at 7-locus MLST genotypes and finally 330-locus cgMLST genotypes. Populations sharing alleles at the 7-locus MLST genotype are referred to as Sequence Types (STs) and members of an ST along with highly related STs (e.g. STs sharing alleles with at least 5 of 7 loci) are considered “clonal complexes”. In *S. enterica*, there are ~360 clonal complexes that are present across 50 of the most common serovars [7].

At the highest levels of resolution (cgMLST), there are hundreds-thousands of different cgMLST genotypes that can be found within a given serotype and within a single ST. Although cgMLSTs belonging to a single ST share broad evolutionary relationships, inferring the hierarchical familial structure of thousands of cgMLST genotypes that may belong to more than one ST is not scalable computationally, particularly when it is necessary to account for horizontal gene transfer (HGT) across divergent lineages. To overcome this problem, multi-locus genotypic classifications can be combined with scalable heuristic Bayesian-based computational approaches such as BAPS, which determines genotypic relationships based on compositional features (admixture) of the core-genome at different scales of resolution. Thus, evolutionary relationships of ST clonal complexes and cgMLST genotypes can be inferred efficiently through hierarchical classification within six BAPS levels (BAPS1 being the lowest level, and BAPS6 the highest level of resolution and population fragmentation). Preliminary analyses showed that multiple STs can be part of the same sub-group within BAPS1, implying they have shared a common ancestor, and this context allows for evolutionary inference of cgMLSTs and their corresponding STs. Hierarchically combining Bayesian admixture-based genotyping schemes at low levels of resolution with ST and cgMLSTs has been shown previously in *Salmonella* [58], as well as other organisms such as *Enterococcus faecium* [59]. Consequently, our heuristic-based approach uses the following hierarchical level of population structure analysis for *Salmonella*: 1) Serovar; 2) BAPS1; 3) STs; and 4) cgMLSTs. It may be noted that, epidemiological clones, which comprise a homogenous population of isolates related to an outbreak, are typically genotyped as cgMLSTs. That happens because cgMLST offers the appropriate level of granularity to define genotypes at the highest level of resolution while considering the shared genomic variation across isolates (i.e. all shared, or >99% loci) [7].

Taxonomically related to *Salmonella* at the Phylum level is the genus *Campylobacter*, which includes two major species (*C. jejuni* and *C. coli*) that are frequent causes of gastrointestinal diseases in humans [60]. *Campylobacter* and *Salmonella* diverge taxonomically at the Class level (*Campylobacter* are members of the Epsilon class of Proteobacteria while *Salmonella* belongs to the Gamma Proteobacteria). Species of *Campylobacter* are also morphologically (helical cells) and physiologically (microaerophilic) distinct from *Salmonella*, but like *Salmonella* can be found in food animals and often are associated with zoonotic outbreaks in developed countries [61].

Though serotypes in *C. jejuni* are defined by combinations of lipopolysaccharide and flagellar antigens, far fewer serotypes are known for *C. jejuni*. However, a distinguishing feature of *C. jejuni* population biology is its propensity for recombination and high frequency of HGT, mediated by specialized systems present in these organisms for uptake and recombination of extracellular DNA [60]. Consequently, *C. jejuni* is less clonal than any given serovar of *S. enterica* lineage I, and it contains a variety of widespread STs, for which the diversification patterns appear to be strongly affected by the host adaptation [60,62]. Consequently, the hierarchical approaches implemented in ProkEvo that can associate genotypes based on BAPS1 admixtures with STs and cgMLSTs provide a powerful means for inferring evolutionary relationships while considering genomic heterogeneity in the clonal frame.

While *Salmonella* and *C. jejuni* are divergent organisms with gram-negative cell wall structures that belong to the phylum Proteobacteria, *S. aureus* is a species that contains a gram-positive cell wall architecture and belongs to the phylum Firmicutes, which is evolutionarily as distinct from Proteobacteia. *Staphylococcus aureus* can cause a diverse array of diseases in humans including skin infections, endocarditis, among others, but it is also a foodborne pathogen that causes foodborne intoxications [63]. Gastroenteritis caused by this pathogen is due to the production of one or more enterotoxins that share some structural features, but can be differentiated serologically. This organism is commonly carried on human skin, nasal cavities, and even the gastrointestinal tract, but also colonizes similar anatomical sites in livestock. In the case of *S. aureus*-associated foodborne illnesses, foods rich in nutrients become contaminated and the organisms produce heat-stable enterotoxins during growth on the food [64]. From WGS data, *Staphylococcus aureus* population can be structured the same way as that of *Salmonella* and *C. jejuni* using BAPS1, STs, and cgMLSTs. However, this pathogen is not as diverse as *C. jejuni* at the ST level, but instead has a degree of clonality that is more comparable to those serovars within *S. enterica* lineage I.

In this era of systems biology and multi-omics methodologies, it is highly desirable to link genotypic classifications of isolates (e.g. serovar, MLST, cgMLST genotypic classifications, and BAPS-based genetic relationships) to important phenotypes associated with resistance to antimicrobial agents, virulence, host adaptation, transmission, and environmental survival. Although linked genotypic and phenotypic data can certainly inform epidemiological surveillance, the linkage affords an even greater opportunity to identify signatures of evolutionary processes (selection) and ecological fitness of the different pathogenic populations in animal and food production environments [95]. Genes and pathways marked by these processes may illuminate selective pressures and better inform risk assessments as well as development of strategies to mitigate spread. Therefore, we designed the studies described below to provide a practical example of how to link the distribution of known AMR genes to the population structure of the organism using serovars and STs in the case of *S. enterica* lineage I serovars, and STs for *C. jejuni* and *S. aureus*. We chose to use known AMR loci and evaluate their association with STs and epidemiological clones worldwide, as in the case of *Salmonella* [65], *C. jejuni* [66], and *S. aureus* [67].

#### Case study 1: *S*. Newport population structure analysis

The *S. enterica* serovar Newport is a zoonotic pathogen that ranks among the top 25 serovars considered as emerging pathogens by the U.S. Centers for Disease Control and Prevention due to several recent outbreaks of foodborne gastroenteritis in humans [96]. Unlike most serovars of *Salmonella enterica* lineage I, which comprise worldwide populations dominated by a single ST clonal complex, the *S*. Newport serovar has diversified into four major STs (*Fig. 4A*). The genetic diversity detected in *S*. Newport isolates is somewhat surprising given its relatively low representation among isolates from the USA available in the NCBI SRA database (total of 2,392 isolates). Thus, this serotype provides a robust example for analysis of a moderately complex population structure through ProkEvo. After the pre-processing steps, assemblies from 2,365 isolates passed the filtering step. The total output data produced by ProkEvo for *S.* Newport was 131GB. After filtering for potentially misclassified genomes using the output of SISTR, 2,317 genomes remained that were annotated as *S.* Newport and predicted as *S*. Newport genotypically (*Fig. S1* and *Fig. S2*). Thus, SISTR-based serovar predictions suggest that 2.03% of the genomes were misclassified as Newport. Using the genotypes assigned by the MLST, cgMLST, and BAPS-based genomic composition programs implemented in ProkEvo, we next defined the relative frequency of each genotype among 2,317 isolates (*Fig. 4A-H*). This analysis identified the expected structure with four dominant STs in the following descending order: ST118, ST45, ST5, and ST132. The cgMLST distribution identified a total of 764 unique cgMLST genotypes, with the cgMLST genotype 1468400426 representing the most frequent lineage or epidemiological clone (*Fig. 4B*) that accounted for ~14% of all isolates whereas the distribution of the other cgMLST genotypes approximately ranges from 0.04 to 4.5%.

**Figure 4:**
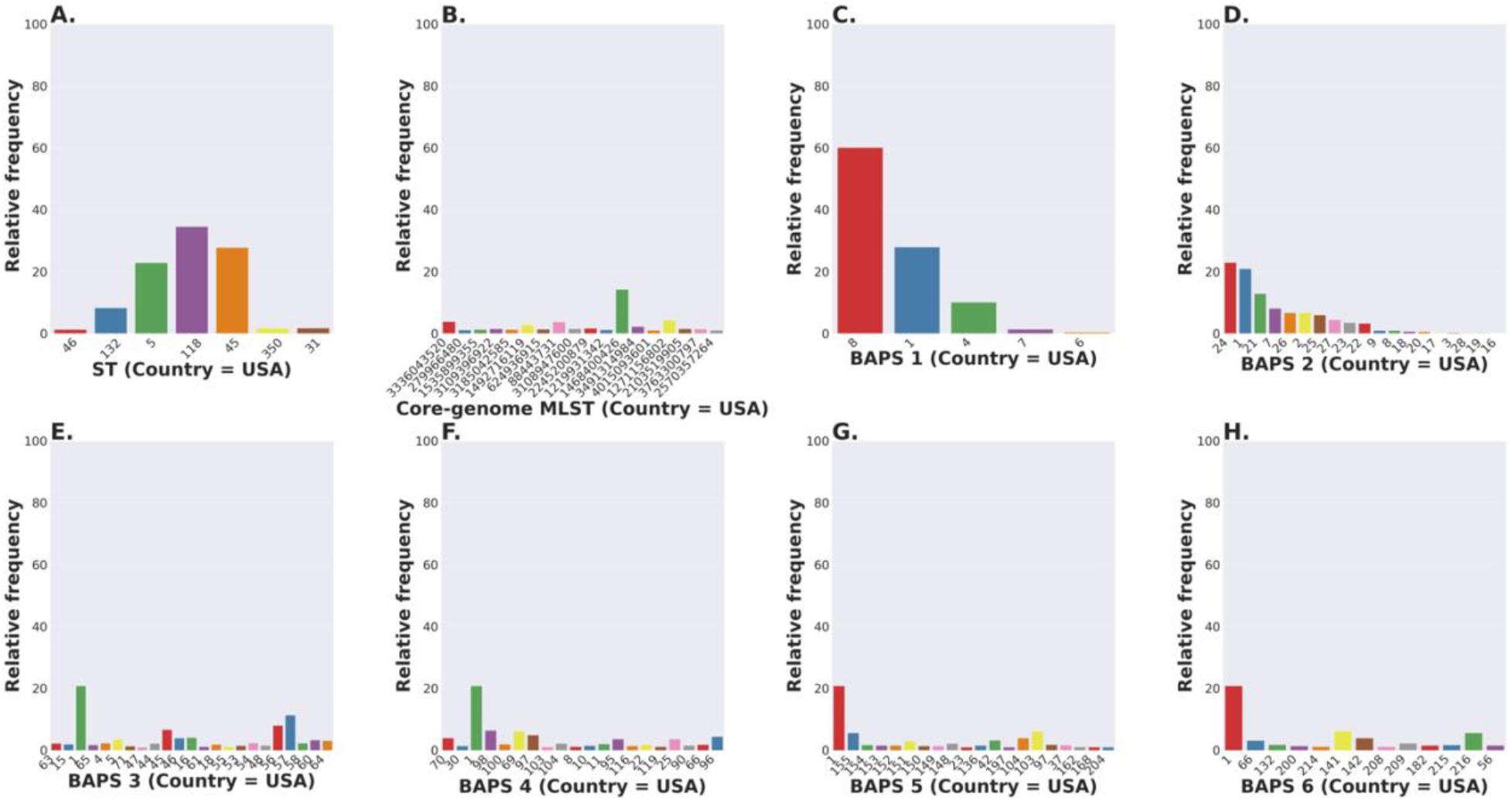
Salmonella Newport (USA) population stratification by genotype classification using two methods: allelic calls (ST and cgMLST) and a heuristic Bayesian approach (BAPS). (A) ST distribution based on seven ubiquitous and genome-scattered loci using the MLST program, which is based on the PubMLST typing schemes (plot excludes STs with relative frequency below 1%). (B) Core-genome MLST distribution based on SISTR which uses ~330 ubiquitous loci (plot excludes STs with relative frequency below 1%). (C-H) BAPS levels 1-6 relative frequencies. For BAPS levels 3-6, we have excluded sub-groups that were below 1% in relative frequency in order to facilitate visualization. Within each BAPS level (1 through 6), each number represents a distinct cluster, or sub-group, to which the isolates belong to. The initial number of genomes used as an input was 2,392, while these analyses were run with 2,365 genomes that passed the post-assembly filtering steps.

To circumvent issues with scalability of phylogenetic inference from thousands of core-genome alignments, we next examined genetic relationships of cgMLST genotypes using the scalable Bayesian-based approach in BAPS to define haplotypes based on the relative degrees of admixture in the core-genome composition at different scales of resolution. As expected, BAPS-based haplotypes at increasing levels of resolution (BAPS1-BAPS6) increasingly fragmented the *S*. Newport into: 9 sub-groups for BAPS1, 32 sub-groups for BAPS2, 83 sub-groups for BAPS3, 142 sub-groups for BAPS4, 233 sub-groups for BAPS5, and 333 sub-groups for BAPS6 (*Fig. 4C-H*). We next used a hierarchical analysis to group the *S*. Newport STs and cgMLSTs based on shared genomic admixtures at BAPS level 1 (BAPS1). At BAPS1, the lowest level of resolution, there are 9 total haplotypes. This analysis showed that the dominant BAPS1 haplotype (BAPS1 sub-group 8) is shared by two of the dominant STs, ST118 and ST5 (*Fig. S1A*). The shared BAPS haplotype implies that the two clonal complexes defined by these dominant STs are more related to each other than to ST45 or ST132, which is consistent with the genetic relationships of these STs predicted by e-BURST [7]. Further analysis of the BAPS1 sub-group 8 haplotype for the major cgMLST lineages also showed 307, 149, and 23 cgMLST genotypes derived from the ST118, ST5, and ST350 clonal complexes, respectively. Having more cgMLST genotypes may suggest that ST118 is a more diverse population, but the diversity may also be inflated compared to other STs based on sample bias and size. Of note, there was not a dominant cgMLST within any of BAPS1 sub-group 8 STs 118, 5, or 350 (*Fig. S1B-D*). Consistent with the genetic relationships of STs predicted by shared BAPS1 sub-group 8 haplotypes, we also found that ST45 belongs to a distinct BAPS1 haplotype (sub-group 1) (*Fig. S1E*), with a total of 152 cgMLST genotypes, and it includes the most frequent cgMLST lineage, cgMLST 1468400426, which is also the most dominant lineage for the entire *S*. Newport dataset (*Fig. S1F*). This predominance of cgMLST 1468400426 within ST45 and across STs, could be due several reasons, including, but not limited to: 1) Sampling effect; 2) Recent outbreaks; 3) Founder effect with a new introduction of a clone in a population; 4) Population drift; or 5) Selective sweep at the whole-genome level in the population due to a selective advantage. Selection and founder effects often underlie the emergence of epidemiological clones that cause significant increases in the numbers of outbreaks [70,71]. While sampling bias obviously contributes to the frequency distribution of genotypes in our analysis, our results are useful to illustrate the types of patterns that can be discovered when using hierarchical population-based analysis, and when used with systematically-designed datasets, that can generate testable hypotheses to define biological features that underlie the frequency distributions of genotypes and populations of interest. In addition to systematic collection of isolates and WGS data, we also emphasize the critical importance of accurate metadata, which is necessary to identify temporal, geographic, or other ecologically-relevant patterns.

After identifying the dominant cgMLST lineage 1468400426, we next examined the degree of genotypic homogeneity in this lineage and compared its genetic relationship to all other cgMLSTs within BAPS1 and ST45 population (*Fig. S2A-E*). We used BAPS-based groupings to estimate genetic relationships of the cgMLSTs to one another. This was accomplished by comparing the frequency of cgMLSTs within BAPS subgroups at increasing levels of BAPS-based partitioning (increasing resolution from level from 2 to 6). To visualize partitioning of the cgMLSTs, genomes belonging to BAPS1 sub-group 1 and ST45 were first selected and then each was categorized into two groups: one group containing cgMLST 1468400426 (numbered 1), and a second group contained all other cgMLSTs (numbered 0). This was done for each genome at each successive level of BAPS2-BAPS6. If the dominant cgMLST 1468400426 is highly clonal, it will be present in one or only a few of the BAPS subgroups at each level of BAPS resolution. As shown in *Fig. S2A-E*, we found that indeed genomes of the dominant cgMLST 1468400426 genotype were all found within a single BAPS subgroup, even at the highest level of resolution (BAPS6). Notably, at each BAPS level, there are other cgMLST genotypes that also co-mapped to the same BAPS subgroups as the dominant cgMLST 1468400426 (sub-group 1 at each level from BAPS2-6 – matching colors between the two stacked bar plots), and the frequency of these other cgMLST genotypes that share BAPS with the dominant cgMLST 1468400426 clone is essentially stable as the BAPS resolution increases. These shared BAPS subgroupings by cgMLST 1468400426 and other co-major lineages including cgMLST 2245200879, cgMLST 843553928, cgMLST 3650140337, cgMLST 4212442350 (in addition to another forty-five minor cgMLSTs) at different levels provide strong evidence of recent common ancestry, and illustrate how admixture analysis such as BAPS can be used to infer evolutionary relationships and circumvent scalability issues with large-scale core genome phylogenies. Importantly, this analysis only defined population stratification within BAPS1 and ST45 clonal complex. Further analysis of cgMLSTs among ST3045, ST3494, ST3783, and ST4493 showed that cgMLST 1468400426 can be rarely found within ST3045 and ST4493, with only one genome of this cgMLST found in each of these two STs. Such a pattern is consistent with the BAPS-based relationships of ST3045, ST3494, ST3783, and ST4493 because these STs all belong to the same BAPS1 sub-group 1 along with ST45.

Collectively, this hierarchical analysis of the genomic relatedness of ST and dominant cgMLST genotypes provides a systematic way to understand population structure and evolutionary relationships of cgMLST genotypes without the need for computationally intensive phylogeny. These relationships are important as they can yield important hypotheses about shared ecological and epidemiological patterns among cgMLSTs that are closely related evolutionarily. It is important to reiterate that this sample of *S*. Newport genomes is from USA. Scaling this analysis to other continents across the globe could reveal if the same population frequencies and inferred evolutionary relationships are observed among global populations. All the steps of these analyses are publicly available in a Jupyter Notebook (https://github.com/npavlovikj/ProkEvo).

#### Case study 2: *S.* Typhimurium population-based analysis

*S.* Typhimurium is the most widespread serovar of *S. enterica* worldwide [97]. Its dominance is partially attributed to its inherited capacity to move across a variety of animal reservoirs including poultry, bovine, swine, plants, and ultimately being capable of infecting humans to cause gastroenteritis or non-Typhoidal Salmonellosis [98,99]. This serovar is phenotypically divided into biphasic and monophasic sub-populations based on their expression of major flagellin proteins from both (biphasic) or only one (monophasic) of the two major flagellin genes [97]. Monophasic *S*. Typhimurium is an emerging zoonotic sub-population and isolates are often resistant to multiple drugs and heavy-metals (copper, arsenic, and silver) [97,100,101]. Due to its relevance as a major zoonotic pathogen and its frequent isolation from clinical and environmental samples, *S.* Typhimurium genomes from a large number of isolates are available (23,045 genomes of isolates from various continents). The large number of genomes available from *S.* Typhimurium dataset is a good measure of the scalability of ProkEvo, since it is an order of magnitude larger than *S*. Newport in the number of genomes. The geographical location of isolates from which these genomes were obtained cannot be determined uniformly from any single field of the associated metadata deposited to NCBI SRA, thus we focus only on scalability, and relationships of STs, cgMLSTs and BAPS-based genotypic partitions to one another. After the download and the pre-processing steps, 21,534 assemblies passed the filtering step, yielding a total data output from ProkEvo of 1.2TB.

As with *S*. Newport, we also conducted an analysis of the population structure based on MLST and cgMLST. However, the sheer size of the *S.* Typhimurium dataset made it necessary to divide the 21,534 genomes into 20 different subsets, with genomes randomly assigned to each subset. Because of this sub-setting, the BAPS-based inquiry of each individual subset would not allow direct comparison of BAPS distributions from each subset because its Bayesian approach.

After quality controlling and filtering the data, we ended up with 20,239 genomes of *S.* Typhimurium biphasic and monophasic combined. In order to present various ways of conducting population-based analyses using ProkEvo, we use combinations of three pieces of information: 1) Whether or not the genome is classified as biphasic or monophasic based on the SISTR algorithm (.csv); 2) The ST clonal complexes calls using the legacy MLST (.csv); and 3) The cgMLST genotypic classification based on SISTR (.csv) [55]. It is important to note that SISTR makes predictions of serotypes based solely on genotypic information. In *Salmonella* that is possible, because of the high degree of linkage disequilibrium between the clonal frame (i.e. genome backbone) and loci that generate the O and H antigens [59]. In the *S.* Typhimurium dataset, 72.6%, 25%, 2.4% of the quality-controlled genomes were classified as Biphasic, Monophasic, or other serovars, respectively. From the Biphasic population, 78.4%, 9.62%, 5.35%, 2.09% of the isolates belonged to ST19, ST313, ST36, and ST34, respectively (*Fig. 5A*). Whereas, for Monophasic, 93%, 5.79%, 0.094% of the isolates belonged to ST34, ST19, and ST36, respectively (*Fig. 5A*). This partitioning matches the known clonality of the *S.* Typhimurium Monophasic populations and their association with the ST34 complex [65]. As for the Biphasic population, it is predominantly associated with ST19 and likely contains the ancestor of the other ST clonal complexes. Most likely, the ST19 dominance is a consequence of its dispersal capacity and ability to spread across a variety of reservoirs, including different species of livestock, and other environments [65]. ST313 has recently emerged in Africa, and is associated with non-Typhoidal Salmonellosis in humans. ST36 represents a minor clonal complex within *S*. Typhimurium Biphasic that appears to be either restricted ecologically, or does not otherwise have fitness advantages to enable it to become widespread, but it is capable of causing gastroenteritis in humans [65].

**Figure 5:**
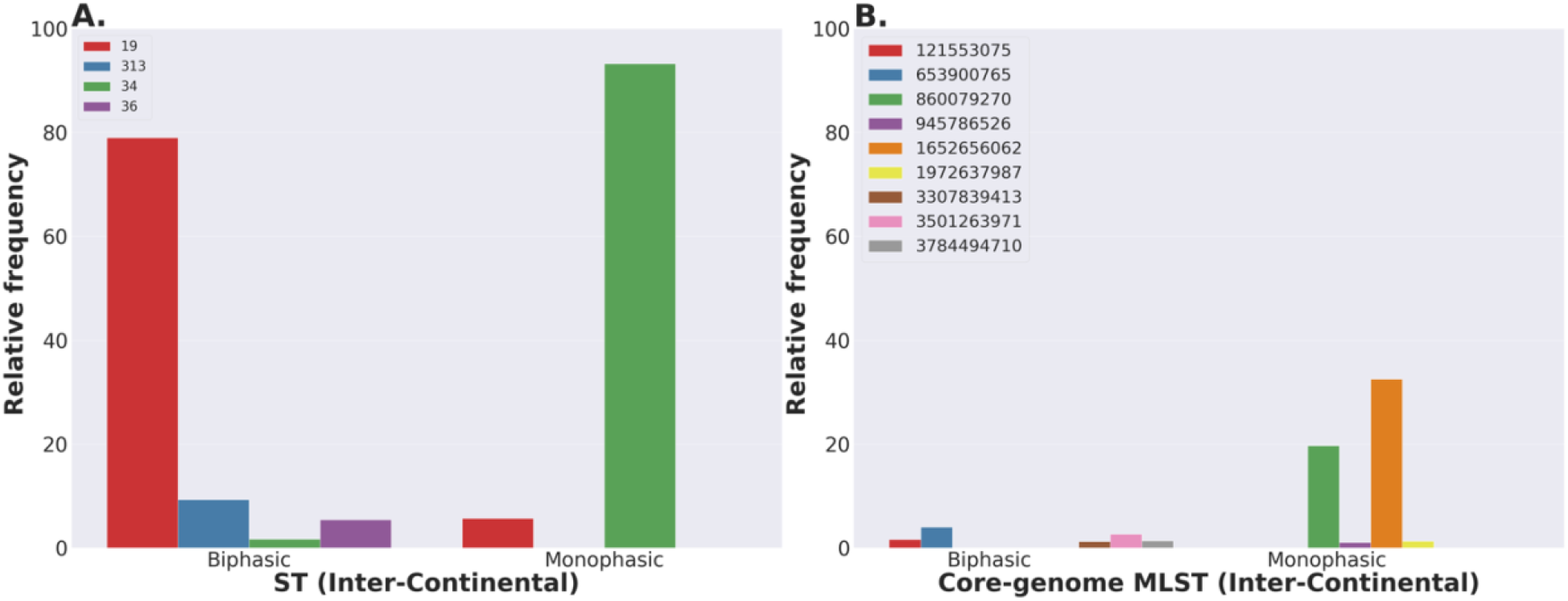
Inter-continental distribution of *Salmonella* Typhimurium STs and core-genome MLSTs. (A-B) Relative frequencies of STs and core-genome MLSTs between Monophasic and Biphasic populations across multiple continents (STs and core-genome MLSTs with proportion below 1% were excluded from the graph). The initial number of genomes used as an input was 23,045, while these analyses were run with 21,534 genomes that passed the filtering steps. Raw sequences were downloaded from NCBI SRA without filtering for USA isolates exclusively. Hence, the name “Inter-Continental”. However, we cannot break the data down into continents, because the metadata was unreliable. Bars that are not visible for a particular ST or cgMLST are either completely absent for that group, or present in a miniscule relative frequency.

In terms of cgMLST genotypic distributions, Biphasic and Monophasic had 5,162 vs. 1,161 unique cgMLST genotypes, respectively. This is expected since the dataset for Biphasic (~75% of the genomes) was larger than Monophasic (~25% of the genomes). Notably, within the Biphasic population there was a large number of cgMLST lineages and each lineage represented only small percentage of the total number of isolates. This pattern would be expected from a diversifying global population that is undergoing random genetic drift and which has not recently experienced a selective sweep. In contrast, the Monophasic population comprised 1,161 total cgMLST lineages, but two cgMLST lineages, cgMLST 1652656062 and cgMLST 860079270, comprised 32.33% and 19.62% respectively, of the total number isolates (*Fig. 5B*). This dramatic difference in frequency distribution of the Monophasic cgMLST lineages could be a consequence of 1) recent selective bottlenecks; 2) Founder effects – new epidemiological clones that have rapidly filled unoccupied ecological niches; 3) higher levels of virulence – isolates are more likely to cause significant disease and more frequent identification of the etiologic agent; 4) isolation bias linked to recent outbreaks; or 5) Population drift. These possible explanations are not exclusive, and they serve to illustrate the importance of systematically using WGS data to monitor frequency distributions of local, regional, national, and global populations and specifically to define when notable changes in frequencies occur as they likely indicate major ecological disturbances or events that otherwise affect populations of the organisms. All the steps for this analysis are shown in our Jupyter Notebook (https://github.com/npavlovikj/ProkEvo).

#### Case study 3: Distribution of known AMR loci across the population structures of *S.* Infantis, *S.* Newport, and *S*. Typhimurium

In case study 3, we illustrate use of ProkEvo to define the distributions of known AMR conferring loci from the Resfinder database across populations of *S. enterica* lineage I (*S*. Infantis, *S*. Newport, and *S*. Typhimurium described above). The goal of this analysis was to show how relationships between the population structures and the distribution of AMR loci can be identified. As described in the Methods section, population-specific results are provided by ProkEvo for several databases such as Resfinder, and the user may specify a database of interest or may elect to use the ProkEvo option of reporting the results comparatively. Our emphasis here on results from Resfinder are driven largely by its broad applications to the fields of ecology and genomic epidemiology [73,74]. Although AMR phenotype predictions based on gene content alone do not have the same precision as measuring AMR phenotypes in the laboratory, monitoring AMR gene frequencies in specific populations of organisms does have the advantage of identifying potentially significant population-scale events that are relevant to public health (e.g. changes in frequencies or new combinations appearing within a population).

Results from the Resfinder analysis identified 72 unique AMR loci in genomes of *S.* Infantis, 125 unique AMR loci in *S.* Newport, and 408 unique AMR loci in *S*. Typhimurium (*Table S1*). All 72 AMR genes in *S*. Infantis were confined to a single clonal complex, ST32. This result was expected because AMR subpopulations of this epidemic clone have emerged in multiple geographies through acquisition of mega-plasmids carrying distinct combinations of AMR genes [68,69]. In contrast, large numbers of AMR loci were found in three of the major clonal complexes of *S.* Newport, with 57 AMR loci in ST118, 84 AMR loci in ST45 and 33 AMR loci in ST5. Similarly, large numbers of AMR loci were found among each of the four dominant clonal complexes of *S*. Typhimirium; ST19 had the most with 301 AMR loci, ST34 had 249 AMR loci, ST36 had 130 AMR loci, and ST313 had 112 AMR loci. Given that ST19 and ST34 are the most frequent clonal complexes in the database for this serovar, it is not surprising that their repertoire of genes would be higher than the others [7,65]. Among the AMR genes identified from any of the three serovars, apparent orthologues of genes known to confer resistance to a broad range of antibiotics were identified, including tetracyclines (*tet* genes), sulfonamides (*sul* genes), macrolides (*mdf* genes), florfenicol and chloramphenicol (*florR* and *catA* genes), trimethoprim (*dfrA* genes), beta-lactamases (*bla* family of genes), and aminoglycosides including streptomycin and spectinomycin (*aph*, *ant*, *aadA*, and *aac* genes) [103].

Although significant numbers of AMR genes were found in many of the dominant STs for each serovar, it should be noted that most of the AMR loci were sparsely distributed among small numbers of isolates within an ST (*Table S1*). Therefore, to visualize population-specific patterns of AMR loci that may be selectively enriched in a lineage, we used a threshold of presence of a given AMR gene in >=25% of the genomes of an individual ST to define the predominant AMR patterns in each population. As illustrated in *Fig. 6*, three major patterns were apparent. First, we note that the patterns of predominant AMR genes were largely somewhat unique to each serovar (*Fig. 6A*). Second, the largest numbers of predominant AMR genes were confined to individual clonal complexes in *S.* Infantis and *S.* Newport and two of the four dominant STs in S. Typhimurium (*Fig. 6B-D*). This may reflect a higher degree of clonality among these STs, but we also point out the potential population homogeneity can be an artefact of oversampling clinical isolates during outbreaks without accounting for the overall environmental diversity. Finally, we note the widespread distribution of the mdf(A)_1 and aac(6’)-Iaa_1 loci across all serovars and all clonal complexes (*Fig. 6A*). Such a high degree of conservation suggests these elements may have been acquired ancestrally, prior to diversification of these three serovars, as opposed to recent independent acquisitions [77]. Also, it is important to mention that we are not differentiating between genes present in chromosome vs. plasmids. The latter are more promiscuous and facilitate HGT between closely related, or divergent populations [104].

**Figure 6:**
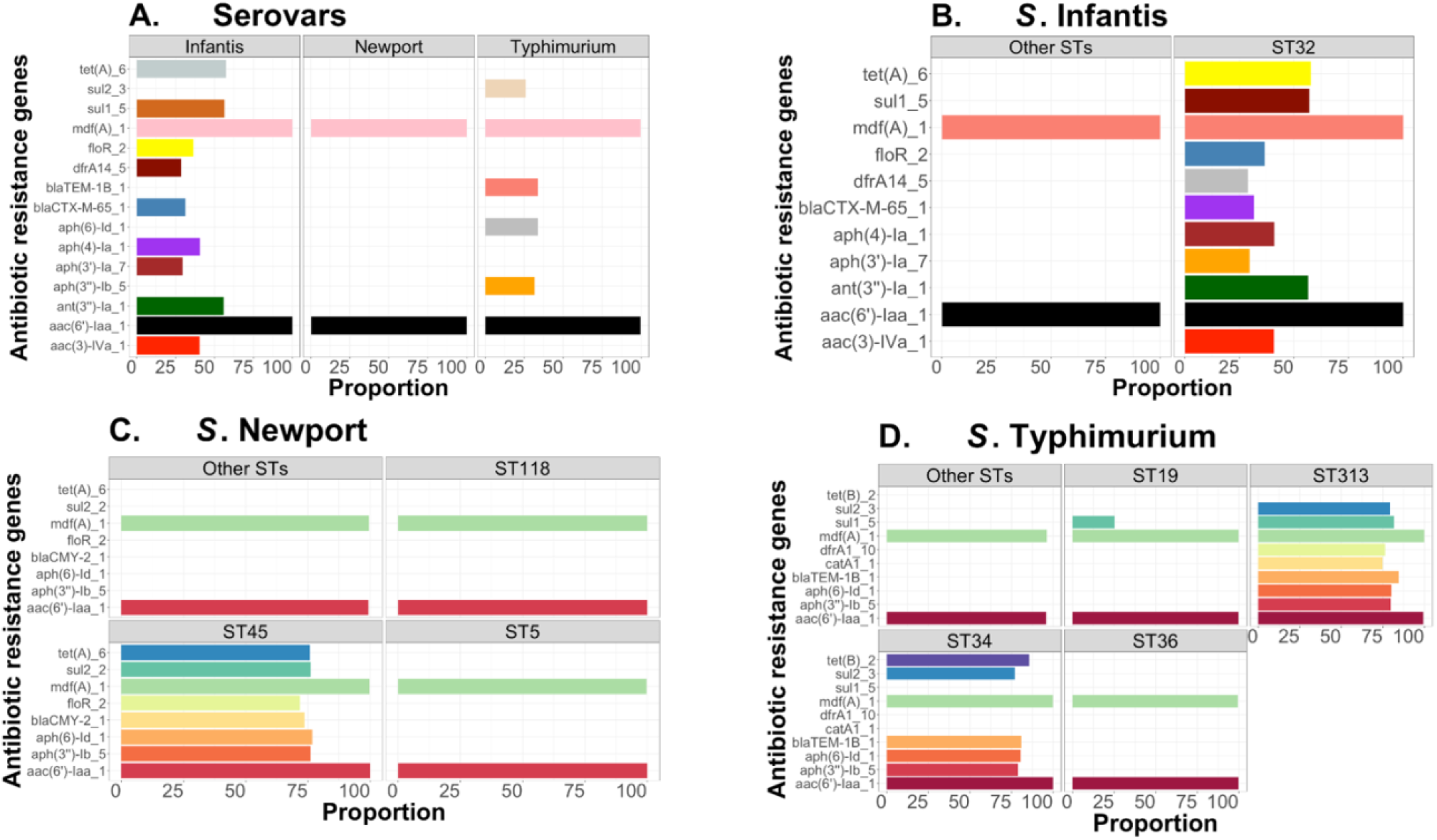
Antibiotic-associated resistance genes distribution between and within three serovars of *S. enterica* lineage I. (A) Proportion of genomes containing antibiotic-associated resistance genes within each serovar. (B-D) Proportion of antibiotic-associated resistance genes within major vs. other STs for *S.* Infantis, *S.* Newport, and *S.* Typhimurium, respectively. For the plots, (B-D), the population was initially aggregated based on the dominant STs vs. the others, prior to calculating the relative frequency of genomes containing each antibiotic-resistance gene. Only proportions equal to or greater than 25% (post-hoc threshold) are shown. For *S.* Infantis and *S.* Newport, only USA data were used; whereas, for *S.* Typhimurium we did not filter based on geography in order to have a larger dataset used to test ProkEvo’s computational performance. Datasets were not filtered for any other epidemiological factor. The total number of genomes used for this analysis was 1,684, 2,365, 21,509 for *S*. Infantis, *S*. Newport, and *S*. Typhimurium, respectively, after filtering out all missing or erroneous values. Also, there were 18 and 1,666 genomes for “Other STs” and ST32 within the *S*. Infantis data, respectively. For *S*. Newport, there were 393, 800, 643, and 529 genomes of the following groups: Other STs, ST118, ST45, and ST5, respectively. Lastly, for *S*. Typhimurium, there were 1,430, 12,477, 1,493, 5,274, and 835 genomes for either Other STs, ST19, ST313, ST34, or ST36, respectively.

#### Case study 4: Population structures and distribution of AMR genes in *C. jejuni* and *S. aureus*

To further illustrate the versatility of ProkEvo across diverse microbial species, we examined the relationships of population structure and AMR gene distributions in two diverse pathogenic bacterial species, *Campylobacter jejuni* and *Staphylococcus aureus*, which belong to very distantly related Phyla. The evolutionary divergence of these taxa can be observed from many unique, fundamental characteristics of *C. jejuni* such as its distinctive morphology (helical) and physiology (microaerophilic). Equally distinctive is the diverse population structure of *C. jejuni*, which features 23 major ST complexes which is largely attributed to unique mechanisms these organisms possess for gene acquisition and recombination, and host adaptation [60,62,81,82].

*Staphyloccus aureus* is a Gram-Positive organism that belongs to the Phylum Firmicutes and is evolutionarily very distantly related to the Proteobacteria. This organism causes an array of infections of the skin and mucosa but can also cause intoxication when it contaminates foods. Its population structure is highly clonal, and three STs (STs 8, 5, and 105) comprise more than 80% of the population (*Fig. 7B*). ST8 is known to be associated with community-acquired infections in the form of either methicillin susceptible or resistant strains (MSSA or MRSA) [84]. ST5 can also cause skin infections and is often found as MRSA [85]; whereas, ST105 is closely related to ST5 and both can carry the *SCCmec* element II [86].

**Figure 7:**
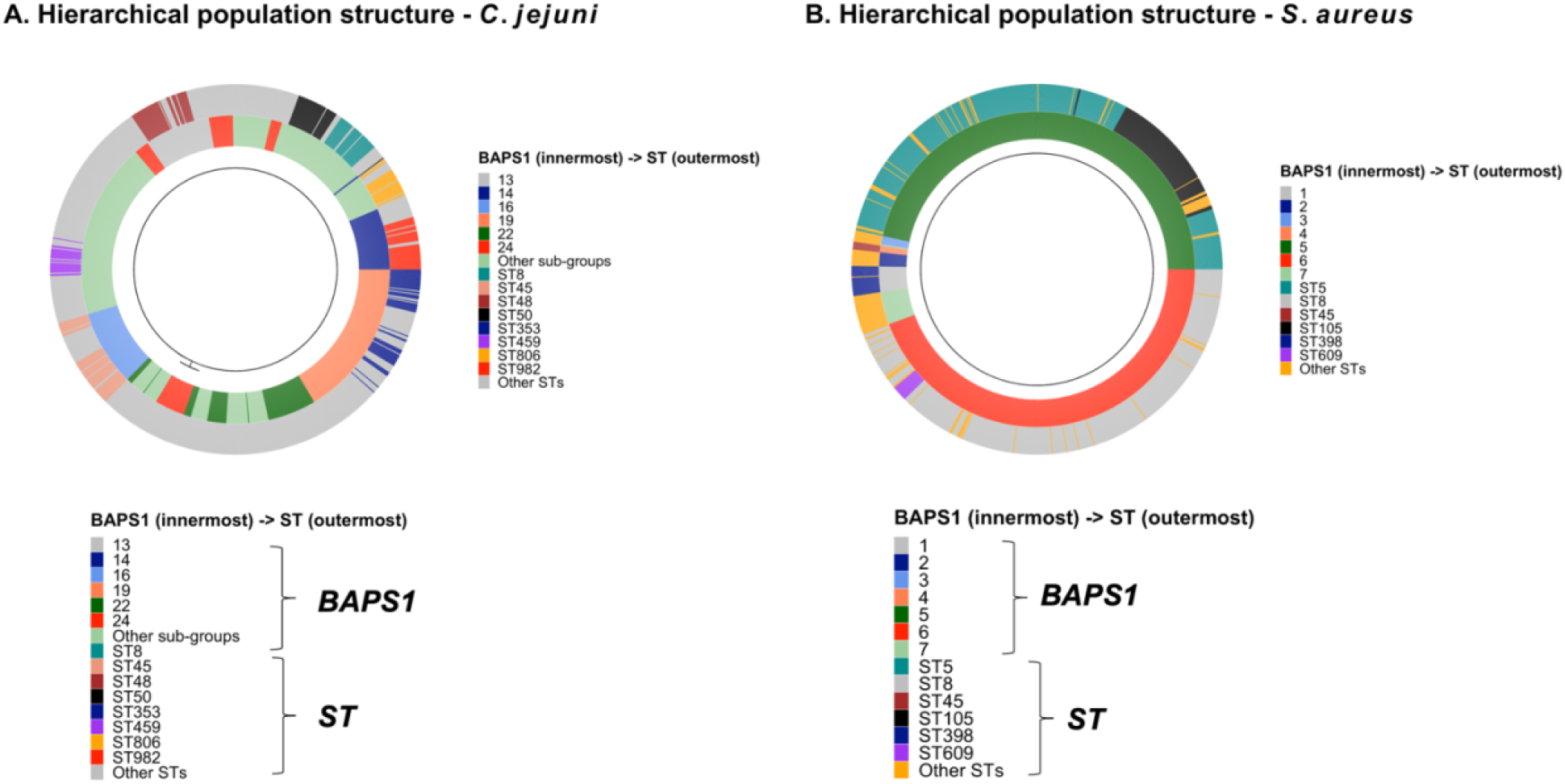
Relationship between the core-genome phylogeny and population structure of *C. jejuni* and *S. aureus*. (A-B) Population structure using BAPS1 and ST for genotypic classifications were overlaid onto the core-genome phylogeny of both *C. jejuni* and *S. aureus*, respectively. BAPS1 was used as the first layer of classification to demonstrate how each sub-group/cluster can be comprised of multiple STs. For instance, STs that cluster together, and belong to the same BAPS1 sub-group, are more likely to have shared a most recent common ancestor. This represents a hierarchical population-based analysis going from BAPS1 to STs. For this analysis and visualization, we have used a random sample composed of 1,044 and 1,193 genomes for *C. jejuni* and *S. aureus*, respectively.

To define hierarchical relationships of major STs from these species to one another, we used the scalable approach described above for relating multi-locus STs to the high-level genotypic groupings from the BAPS level 1 admixture analysis from ProkEvo. In addition to the bar charts, we also illustrate an integrative approach to visualize the frequencies and inter-relationships of STs and BAPS-based genotypes (*Fig. 7A-B*), where each genome is depicted as a member of a circular graph, and their coloration in concentric rings depicts their associated BAPS genotypes (innermost ring) and ST genotypes for each genome. As shown in *Fig. 7A*, this visualization illustrates how genomes from individual ST complexes of *C. jejuni* are found within a single BAPS1 genotype. For example, the ST45 clonal complex is found exclusively within the BAPS1 sub-group 16, ST48 is confined within the BAPS1 sub-group 13, ST353 is found within the BAPS1 sub-group 19, and ST982 is found within the BAPS1 sub-group 14. Thus, despite the extensive contribution of HGT and recombination to genetic diversity in *C. jejuni*, the scalable, hierarchical combination of Bayesian-based admixture analysis and multi-locus genotyping in ProkEvo still enables detection and visualization of broad evolutionary relationships of the STs to one another. Similarly, visualization of the relationships between BAPS1-level genotypes and the most frequent STs in *S. aureus* (*Fig. 7B*) illustrated the high-degree of clonality in its population structure. The dominant ST5 and ST105 were found exclusively within the BAPS1 sub-group 5, ST398 was restricted to the BAPS1 sub-group 1, and the ST609 complex was found within the BAPS1 sub-group 6.

Using the framework of hierarchical BAPS-ST relationships, we next used ProkEvo outputs from the Resfinder database to examine distributions of AMR genes among the STs in these diverse organisms. For this analysis, we focused on STs representing >1% of the total number of genomes for both *C. jejuni* and *S. aureus* (*Fig. 8A-B*). The ProkEvo-mediated search of the Resfinder database from *C. jejuni* genomes identified 256 unique AMR elements in *C. jejuni* and 164 AMR loci for *S. aureus*. Within *C. jejuni*, the top 8 most frequent STs had the following total number of AMR loci: ST353 (29), ST45 (30), ST982 (20), ST48 (24), ST50 (31), ST8 (20),

**Figure 8.**
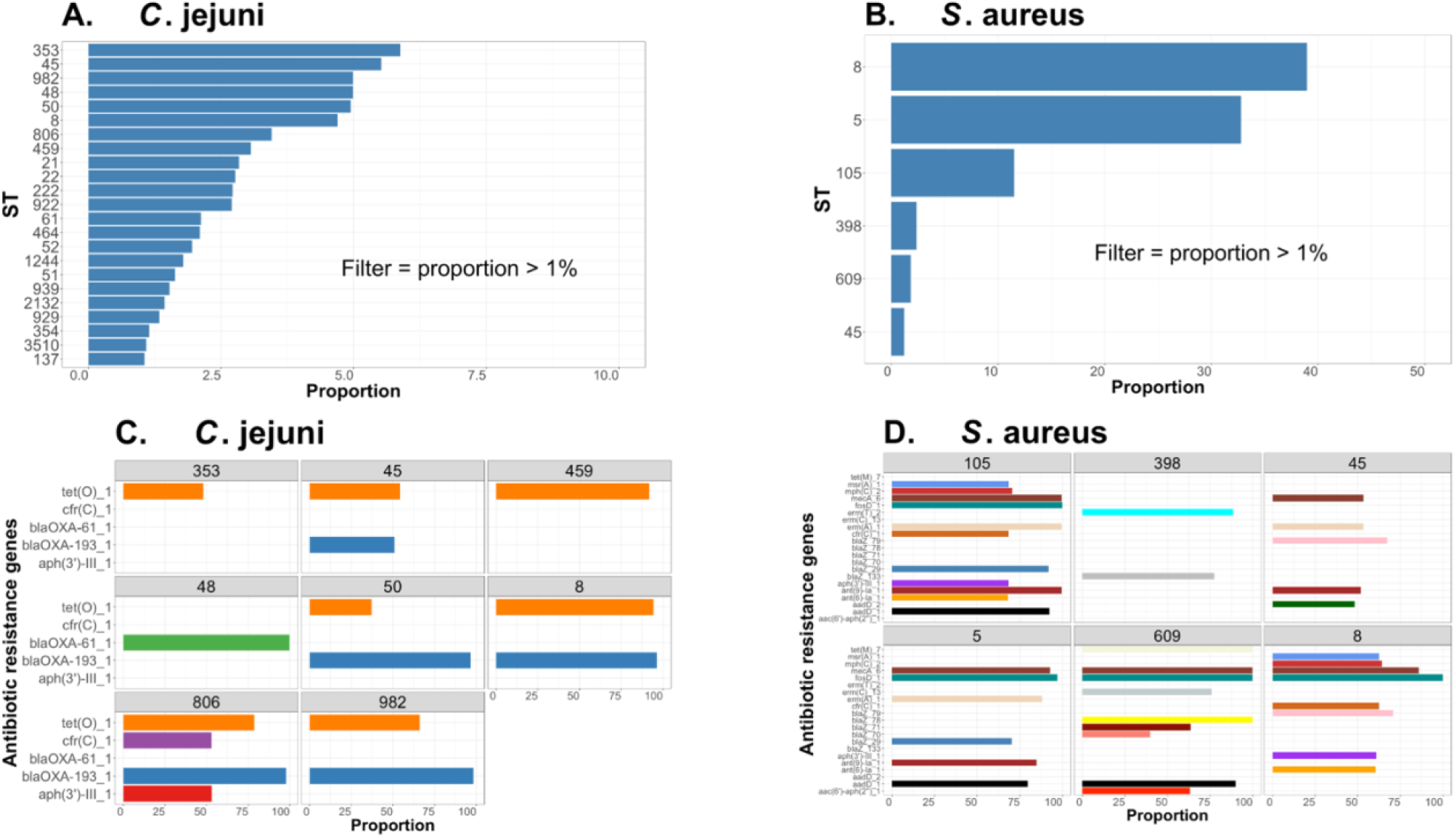
ST-based population structure and distribution of antibiotic-associated resistance genes for two major foodborne pathogens. (A-B) Proportion of the most dominant STs within *C. jejuni* and *S. aureus* populations (only proportions >1% are shown). (C-D) Proportion of genomes containing antibiotic-resistance genes within ST populations for *C. jejuni* and *S. aureus* (only proportions >25% are shown). Both datasets only included genomes from USA and were not filtered for any other epidemiological factor. The total number of genomes entered in this analysis was 18,845 and 11,597, for *C. jejuni* and *S. aureus*, respectively, after filtering out all missing or erroneous values. For *C. jejuni*, there were 886, 1,041, 940, 932, 1,108, 577, 651, and 940 genomes of the following groups: ST8, ST45, ST48, ST50, ST353, ST459, ST806, and ST982, respectively. Lastly, for *S. aureus*, there were 4,518, 3,801, 1,334, 276, 211, and 141 genomes for either ST8, ST5, ST105, ST398, ST609, or ST45, respectively.

ST806 (19), and ST459 (15). Thus, there was relatively even distribution of AMR loci among the most dominant STs. In contrast, the number of AMR loci in *S. aureus* was essentially a function of the frequency of the STs. The most frequent *S. aureus* STs in the database (ST8, ST5, and ST105) contained the largest number of AMR loci (ST8 had 88 AMR loci, ST5 had 85 AMR loci, and ST105 had 52 AMR loci). In contrast, the lower frequency STs had fewer (ST398 had 39 AMR loci, ST609 had 20 AMR loci, and ST45 had 24 AMR loci). The link to the intermediate files used to obtain this information can be found in *Table S1*.

As was the case with distributions of AMR loci in *Salmonella enterica*, most of the AMR loci detected in *C. jejuni* and *S. aureus* were sparsely distributed across isolates belonging to an individual ST. Therefore, we focused on AMR loci in >=25% of the isolates within an ST. As shown in *Fig. 8*, despite the fact that *C. jejuni* had a greater number of total AMR-associated genes, *S. aureus* had a greater number of prominent AMR loci meeting the >25% threshold.

Although the data are certainly biased to clinical samples, it is tempting to speculate that selection for AMR in *S. aureus*, which is most often transmitted from human-human, is likely much stronger than zoonotic *Campylobacter jejuni*. The diversity of prominent AMR loci in *S. aureus* was also quite noticeable, with each of the major STs having a distinct combination of AMR loci.

With respect to *C. jejuni*, there was widespread co-distribution of tet(O)_1 and blaOXA-193_1, which confer resistance to tetracyclines and beta-lactamases, respectively, across the five most frequent STs (*Fig. 8C*). One potential explanation for this pattern would be an ancestral acquisition of both genetic elements, and subsequent loss of one or both genes during divergence of the ST48, ST353, and ST459 populations [87]. In contrast, the *cfr(C)_1* and *aph*(3’)-III_ loci appear uniquely in the *C. jejuni* ST806 clonal complex, suggesting these genes are relatively recent acquisitions within the clonal complex. The *cfr* gene is of great interest because it has a pleiotropic phenotype associated with resistance to a variety of AMR classes, such as: phenicol, lincosamide, oxazolidinone, pleuromutilin, streptogramin A, and other macrolides [88].

In the case of *S. aureus*, the largest number of prominent AMR genes were found in STs 5 and ST105 (*Fig. 8D*), both of which belong to the same BAPS1 sub-group 5 genomic type, and are thus more closely related to each other than the other dominant STs (*Fig. 7B*). ST8 and ST609 also carry significant numbers of prominent AMR loci and these STs also share evolutionary history, since they both belong to BAPS1 sub-group 6 genomic type (*Fig. 7B*). In contrast, ST398 and ST45 contain the fewest AMR loci and each belongs to a distinct BAPS1 sub-group (ST398 is a member of BAPS1 sub-group 1 while ST45 is a member of BAPS1 sub-group 4 – *Fig. 7B*).

## Discussion

The continuous increase in the volume of WGS data from bacterial species is driving the field of bacterial genomics away from simple comparative and functional genomics towards population-scale inquiry. This shift requires approaches rooted in data science to process, analyze, and mine WGS data at scales that have not before been achieved. Indeed, the vast number of genomes currently available is already driving development of tools, pipelines, and approaches for analysis of population dynamics, phylogeography, and epidemiological patterns at whole genome-scales of resolution. When scalable tools for population-based inquiry at genomic resolution are combined with appropriate sampling of environments and robust metadata, these unique approaches will collectively provide entirely new ways to understand fundamental ecology of important microorganisms, environmental factors that drive ecological adaptation, and the evolutionary mechanisms through which such adaptations are mediated [3,4,7,10,16].

Scalability remains one of the key bottlenecks that limits population-based inquiry at genomic resolution. Automation and parallelization of complex pipelines for implementation on different types of computational platforms (e.g. clusters and grids) can help overcome the scalability bottleneck. ProkEvo fills this gap by allowing researchers to scale and automate the analyses from hundreds to many thousands of genomes without the need to write individual scripts to run programs and move data input/output from program to program. Indeed, such approaches become difficult to reproduce across laboratories. ProkEvo takes advantage of a set of well-developed bioinformatics tools and a robust workflow management system that enables implementation on different high-performance and high-throughput computational platforms. Thus, ProkEvo produces scalable and highly reproducible workflows. We acknowledge that full automation of workflows in ProkEvo has trade-offs, because users may rely on such systems without understanding how the underlying assumptions and tuning of important parameters in the individual programs ultimately affect their studies. On the other hand, a large number of biology/microbiology laboratories can immediately benefit from the automation and scalability of ProkEvo to generate a variety of novel hypotheses. Moreover, implementation of ProkEvo in these research environments will ultimately become a major force to drive development of more systematic approaches to study designs for large-scale studies and ongoing surveillance, including the sampling collection of WGS from isolates, as well as the collection and curation of critical metadata.

ProkEvo is modular, and each genome is analyzed independently when computing resources are available. In theory, if a dataset has *n* genomes and a computational platform has *n* available cores, ProkEvo can easily scale linearly and utilize all resources at the same time on execution platforms such as clusters and grids. ProkEvo only needs a list of NCBI SRA (genome) identifications as an input, and the Pegasus submit script. The computational resources used for the steps in ProkEvo are specified per tool and are not fixed. This is an important feature of ProkEvo that allows efficient allocation of resources and requires high resources only when needed. While the scripts for executing the tools in ProkEvo are written to consider common errors such as bad input data or exceptions, failures due to rare cases are still possible. In such instances, only the failed job is retried, with the possibility of terminating only the failed job upon repeated failure. Failure of individual jobs does not affect the continuity of the pipeline and the remaining independent jobs continue running. This feature is extremely useful when analyzing large datasets, and bypasses the problem of very small fractions of the tens of thousands of genomes having faulty reads that would otherwise disrupt the entire workflow across all jobs.

Most of the enabling capability of ProkEvo relies upon automation and management of the massive workflows through Pegasus WMS. The scalability, ability to handle large sets of data with complex input-output dependencies, and portability to different computational platforms are just few of the advantages that drove selection of Pegasus as the WMS for ProkEvo. Although we used ProkEvo in this report to efficiently process and analyze a moderately large dataset of >20,000 genomes from *Salmonella* Typhimurium, future testing needs to be done to evaluate and improve ProkEvo’s performance with hundreds of thousands of genomes. Additionally, its portability to cloud environments such as the Amazon Web Service needs to be evaluated.

Despite the efficiency of the Pegasus WMS, one of the central programs of ProkEvo (Roary) creates a bottleneck in generating core-genome alignments. This step is important since it precedes population structure analysis using fastbaps or downstream phylogeny, and it can run indefinitely when the number of genomes is large. Our workaround here was to randomly divide the dataset into subsets of up to 2,000 genomes, which allows ProkEvo to perform all jobs efficiently. However, this approach has consequences because: 1) fastbaps uses Bayesian BAPS computations which may confound direct data aggregation afterwards; 2) the user will have to generate multiple phylogenetic trees; and 3) pan-genome annotation may vary across subsets and there may be inconsistent gene calls/classifications, particularly with respect to hypothetical proteins. We are examining other scalable computational approaches for phylogenetic inference such as kmer-based construction of distance matrices using raw reads directly [105]. An advantage of ProkEvo is that the Pegasus WMS can easily accommodate addition of novel programs/algorithms to the platform without disrupting any pre-established tasks. Hence, new solutions or alternative steps or programs can easily be incorporated into the ProkEvo workflow.

The versatile Pegasus WMS has been used for development of small and large-scale processing and computational pipelines for a variety of projects and applications across multiple disciplines, including LIGO gravitational wave detection analysis [51], the structural protein-ligand interactome (SPLINTER) project [90], the Soybean Knowledge Base (SOyKB) pipeline [91], and the Montage project for science-grade mosaics of the sky [92]. To the best of our knowledge, use of an advanced WMS such as the Pegasus WMS is a very unique feature of ProkEvo that is not found in other complex pipelines for large-scale analysis bacterial genomes such as EnteroBase [17], TORMES [18], Nullarbor [19], and ASA3P [20]. EnteroBase is an online resource for identifying and visualizing bacterial species-specific genotypes by utilizing a high-performance cluster at the University of Warwick. TORMES is a whole bacterial genome sequence analysis pipeline that works with raw Illumina paired-end reads, and is written in Bash. Nullarbor is a Perl pipeline for performing analyses and generating web reports from sequenced genomes of bacterial isolates for public health microbiology laboratories. ASA3P is an automated and scalable assembly annotation and analyses pipeline for bacterial genomes written in Groovy. In addition to the Pegasus WMS, ProkEvo is also distinct from these programs in its ability to combine classifications of each genome based on multi-locus genotypes (at ST and cgMLST scales) with the scalable approach of classifying genomic types based on Bayesian admixture analysis.

As illustrated in all four case studies, our approach of combining hierarchical combinations of genotypic classifications with phenotypes such as AMR can produce novel insights. In these case studies, we demonstrated how the combination of multi-locus approaches and Bayesian admixture analysis can illuminate evolutionary relationships with scalable methods. Our studies also identified combinations of AMR genes that are widely dispersed across dominant clones as well as AMR genes with population-specific patterns of distribution. Integrating population genomics (allele and clone frequencies) outputs from ProkEvo with complex trait analyses can begin to identify casual variants that are driving evolution and ecological characteristics [109]. Population-based selective sweeps (i.e. purged genomic variation at the whole genome level) can be driven by acquisition of a single locus capable of providing novel physiological or virulence trait [77], as exemplified by acquisition of novel loci in clonal complexes ST21 and ST45 of *C. jejuni* which reduce oxygen sensitivity and enhance survival and spread across the poultry food chain [10]. However, complex traits such as ecological fitness can also arise from contributions of allelic variation at multiple loci [3,78]. Of course, this is not a one-way street as such variation in bacterial pathogens is also met with variation in host loci that contributes to susceptibility, for instance [79].

It is important to note that our case studies drew upon the large numbers of genomes already available in the NCBI SRA databases, and we have been careful to draw only general conclusions because of the inherent bias in broad species or serovar-specific datasets. The most common bias in WGS representations of pathogenic species results from overrepresentation of clinical samples in general and potential oversampling large numbers of isolates from outbreaks and epidemiological clones. Such representation does provide important temporal approaches for detecting frequency changes in dominant clones, which even at high-levels of resolution are indicative of significant changes in transmission patterns. However, to truly understand the ecology of these populations and how their ecological characteristics in livestock and the environment relate to transmission and virulence will require systematic sampling and accurate estimates of clone frequency from those environments for robust comparison to those found in clinical samples. Epidemiological lineages (i.e. cgMLST clones) making all the way to human clinical cases are by definition “successful”. However, the cgMLST distribution, and pattern of dominance, could have arisen by random chance (i.e. neutral selection), and subsequently be maintained by the influence of habitats that facilitate the survival and spread of a given clone(s) [110]. Importantly, the ProkEvo platform should facilitate systematic collaboration and coupling of analysis from ongoing surveillance studies and regulatory testing in animal and food production environments.

In addition to bias in sample types, our analyses were also limited by the availability of standardized formats for associating metadata with WGS data in the SRA. Even the most basic type of information such as isolation date is not uniformly available or is not consistently entered into the same fields, which is required for automation. Ongoing efforts from consortia led by NIST and other agencies are making progress, but significant barriers to data sharing across regulatory, industry, and academic sectors still exist [80]. However, we believe the opportunity for developing entirely new approaches to mitigation and control of pathogenic bacteria that will result from fundamental understanding of their ecology and evolution across entire food production systems will ultimately outweigh the risks of making such data and metadata publicly available. Indeed, experimental designs that incorporate systematic sampling and standardized metadata can then be coupled with modern statistical tools such as machine learning and pattern searching algorithms [106,107,108] that can be easily implemented in ProkEvo. Such approaches will enable communities of microbiologists, epidemiologists, and bacterial geneticists across the academic, regulatory, and industrial sectors to truly exploit the massive amount of emerging WGS data.

## Conclusions

In this paper we describe the **ProkEvo platform**, which is: 1) An automated, user-friendly, reproducible, and open-source platform for bacterial population genomics analyses that uses the Pegasus Workflow Management System; 2) A platform that can scale the analysis from at least a few to tens of thousands of bacterial genomes using high-performance and high-throughput computational resources; 3) An easily modifiable and expandable platform that can accommodate additional steps, custom scripts and software, user databases, and species-specific data; 4) A modular platform that can run many thousands of analyses concurrently, if the resources are available; 5) A platform for which the memory and run time allocations are specified per job, and automatically increases its memory in the next retry; and 6) A platform that is distributed with conda environment and Docker image for all bioinformatics tools and databases needed to perform population genomics analyses. Our case studies illustrate how to perform an initial population-based analyses using ProkEvo output files with reproducible Jupyter Notebooks and R scripts. Results from our case studies are clear illustrations of the types of evolutionary and ecological inquiry that can be made from large-scale, WGS-based datasets using the ProkEvo platform.

## Acknowledgements

This work was completed by utilizing the Holland Computing Center of the University of Nebraska, which receives support from the Nebraska Research Initiative, and using resources provided by the Open Science Grid, which is supported by the National Science Foundation and the U.S. Department of Energy's Office of Science. We would like to greatly thank Mats Rynge for his extensive assistance and valuable suggestions while setting up and running ProkEvo on the Open Science Grid. We also thank Dr. Derek Weitzel and Karan Vahi for their technical support.

This paper is dedicated to the memory of Dr. David Swanson, the former director of the Holland Computing Center, who passed away before this project was completed. Dr. David Swanson was an amazing individual, a true leader and an inspiration to us all. He was a strong advocate of computational research and literacy, and passionate about finding ways to do large-scale science attainably, better, and faster. Our collaboration and shared endeavors would not have been possible without his support and guidance.

