## Supplemental Files for "ProkEvo: an automated, reproducible, and scalable framework for high-throughput bacterial population genomics analyses"

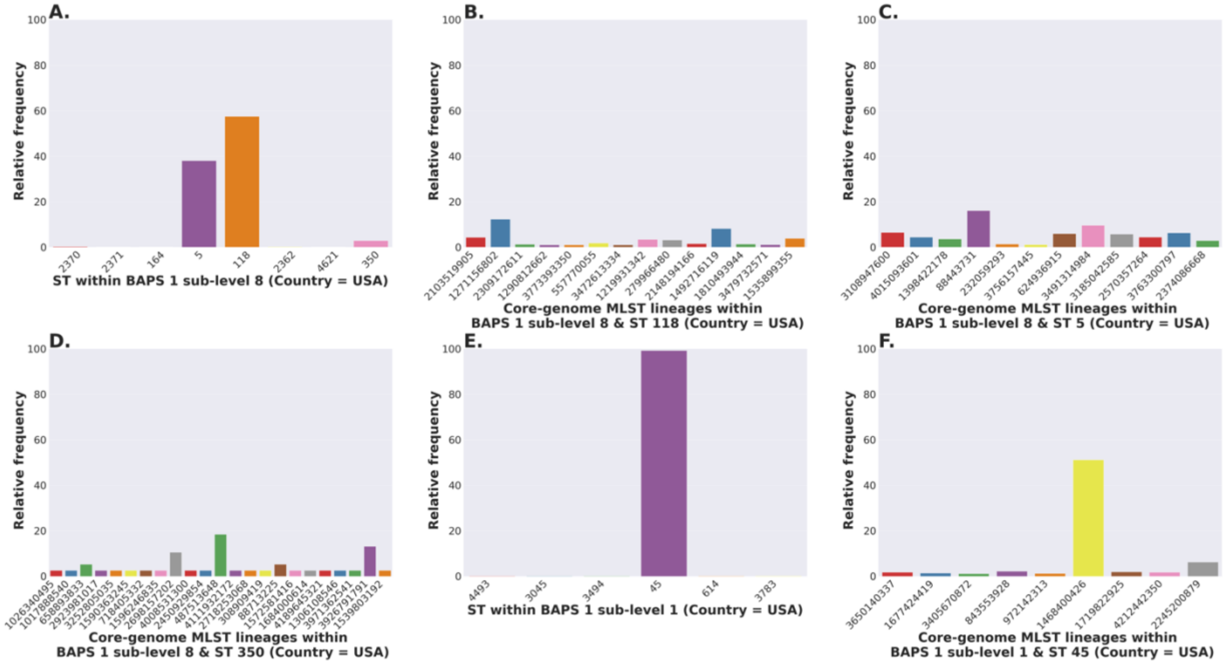


**Figure S1: *Salmonella* Newport (USA) hierarchical population-based analysis.**
(A) ST distribution with BAPS1 sub-group (equals sub-level) 8 (excluding STs with relative frequency below 0.1%). (B) Core-genome MLST lineage distribution within BAPS1 sub-group 8 and ST118 (excluding lineages with relative frequency below 1%). (C) Core-genome MLST lineage distribution within BAPS1 sub-group 8 and ST5 (excluding lineages with relative frequency below 1%). (D) Core-genome MLST lineage distribution within BAPS1 sub-group 8 and ST350 (excluding lineages with relative frequency below 1%). (E) ST distribution with BAPS1 sub-group 1 (excluding STs with relative frequency below 0.1%). (F) Core-genome MLST lineage distribution within BAPS level 1 sub-level 1 and ST45 (excluding lineages with relative frequency below 1%). The number of filtered genomes (i.e. genomes that passed assembly quality control metrics) used as an input in this analysis was 2,365.

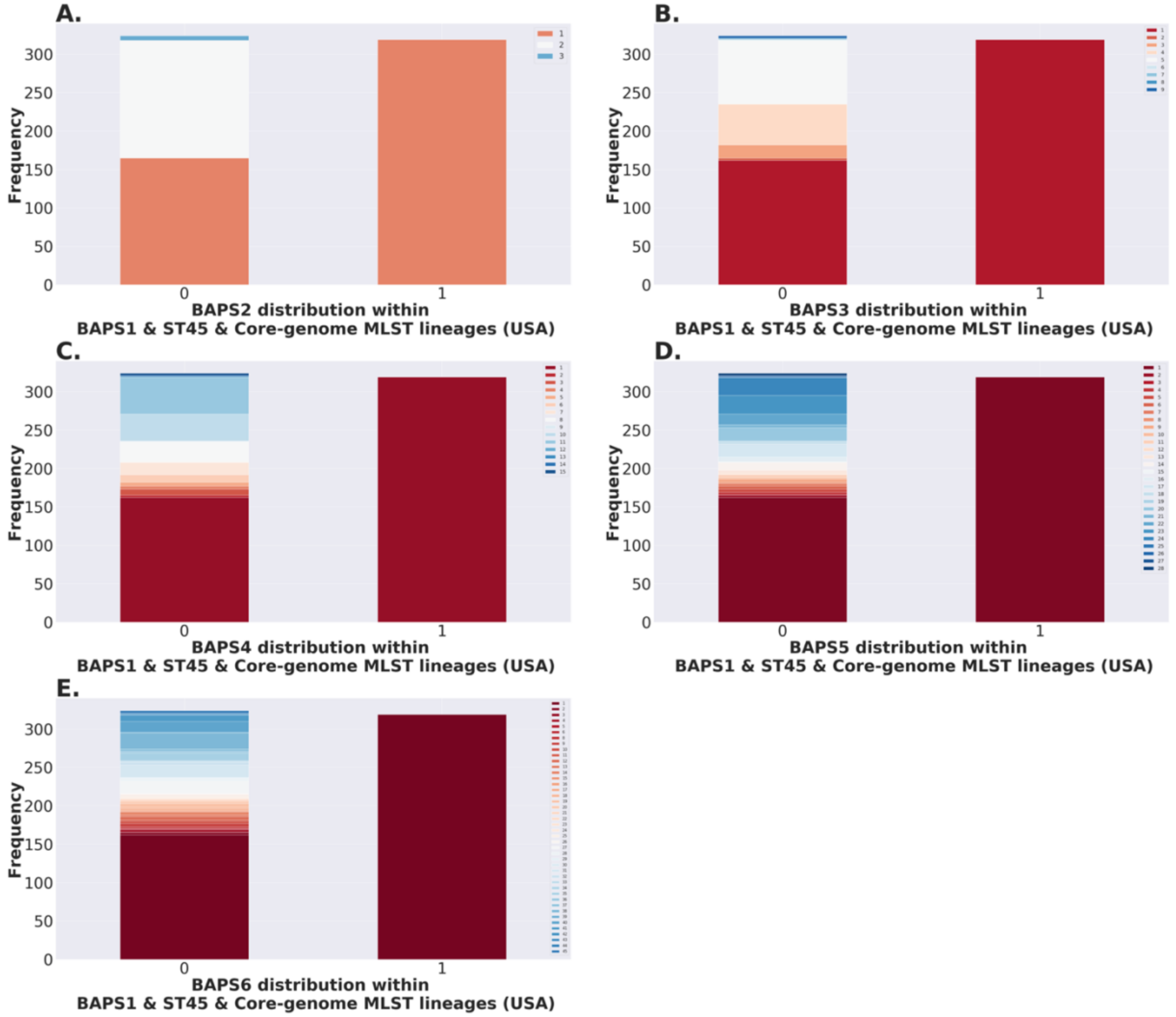


**Figure S2: BAPS levels 2-6 frequency of core-genome MLST lineages with BAPS sub-group 1 and ST45 within *S.* Newport.**

First, core genome MLST (cgMLST) genotypes were classified as 1 if it were cgMLST 1468400426, and 0 otherwise. cgMLST 1468400426 can be called a major epidemiological clone (i.e. higher relative frequency) within BAPS1 sub-group 1 and ST45. The goal was to compare the distribution of BAPS levels 2-6 between cgMLST 1468400426 or the other epidemiological clones as a sub-population of BAPS1 sub-group 1 and ST45. (A-E) Frequency of BAPS levels 2-6, respectively, when comparing group classified as 1 (core genome MLST = 1468400426) or 0 (core genome MLST = others), within BAPS sub-group 1 and ST45, as part of a hierarchical approach for analysis of the *S.* Newport population in the United States (USA). Specifically, with this analysis one can evaluate how genotypically homogenous (i.e. clonal) the population of that epidemiological clone is when compared to other cgMLSTs all combined. A highly clonal population will have few or even a single BAPS sub-group as the levels go up from BAPS2 to BAPS6. A more diverse epidemiological clone will have the number of sub-groups increased, the more one stratifies the population going from BAPS2 to BAPS6. The number of quality-controlled genomes used as an input for these analyses was 2,365.

**Table S1: Location of the intermediate files used to generated the Figures in this paper. While the raw files are available on FigShare, the intermediate files include additional filtering used to produce the corresponding Figures.**

| | **Link** |
|---|---|
| **Distribution of antibiotic resistance genes across STs of *S.* Infantis** | https://figshare.com/articles/dataset/abx_serovars_infantis_csv/13082906 |
| **Distribution of antibiotic resistance genes across STs of *S.* Newport** | https://figshare.com/articles/dataset/abx_serovars_newport_csv/13083032 |
| **Distribution of antibiotic resistance genes across STs of *S.* Typhimurium** | https://figshare.com/articles/dataset/abx_serovars_typhimurium_csv/13083176 |
| **Distribution of antibiotic resistance genes across 3 serovars of *S. enterica* lineage I** | https://figshare.com/articles/dataset/abx_serovars_salmonella_csv/13082795 |
| **Relative frequencies for *C. jejuni* STs** | https://figshare.com/articles/dataset/c_jejuni_st_distribution_csv/13082771 |
| **Antibiotic resistance gene distribution across major STs of *C. jejuni*** | https://figshare.com/articles/dataset/abx_major_c_jejuni_st_csv/13082777 |
| **Relative frequencies for S. *aureus* STs** | https://figshare.com/articles/dataset/saureus_st_distribution_csv/13082783 |

| | |
|---|---|
| **Antibiotic resistance gene distribution across major STs of S. *aureus*** | https://figshare.com/articles/dataset/abx_major_saureus_st_csv/13082786 |
